# Group B Streptococcus Adaptation Promotes Survival in a Hyperinflammatory Diabetic Wound Environment

**DOI:** 10.1101/2022.06.27.497804

**Authors:** Rebecca A. Keogh, Amanda L. Haeberle, Christophe J. Langouët-Astrié, Jeffrey S. Kavanaugh, Eric P. Schmidt, Garrett D. Moore, Alexander R. Horswill, Kelly S. Doran

**Affiliations:** Department of Immunology and Microbiology, University of Colorado Anschutz, Aurora, CO, USA; Department of Medicine-Pulmonary Sciences and Critical Care, University of Colorado Anschutz, Aurora, CO, USA; Department of Veterans Affairs Eastern Colorado Healthcare System, Aurora, Colorado, USA; Department of Orthopedics, University of Colorado, Anschutz Medical Campus, Aurora, CO, USA

## Abstract

Diabetic wounds have poor healing outcomes due to the presence of numerous pathogens and a dysregulated immune response. Group B *Streptococcus* (GBS) is commonly isolated from diabetic wound infections, but the mechanisms of GBS virulence during these infections have not been investigated. Here, we develop a murine model of GBS diabetic wound infection, and using dual RNA-sequencing, demonstrate that GBS infection triggers an inflammatory response. GBS adapts to this hyperinflammatory environment by upregulating virulence factors including those known to be regulated by the two-component system *covRS*, such as the surface protein *pbsP*, and the *cyl* operon which is responsible for hemolysin/pigmentation production. We recover hyperpigmented/hemolytic GBS colonies from the murine diabetic wound which we determined encode mutations in *covR*. We further demonstrate that GBS mutants in *cylE* and *pbsP* are attenuated in the diabetic wound. This foundational study provides insight into the pathogenesis of GBS diabetic wound infections.

**Teaser:** The Fight for Survival by the Bacterium Group B Streptococcus in the Diabetic Wound.

## Introduction

Diabetes is a multifaceted metabolic disease that is estimated to affect more than 500 million individuals worldwide by 2035 (1). One of the most common complications in diabetic individuals is the development of wounds such as foot ulcers, which an estimated 19-34% of all patients will develop in their lifetime (2). Normal wound healing progresses in four overlapping stages; (i) hemostasis, (ii) inflammation, (iii) proliferation and (iv) remodeling (3). However, diabetic wounds are often stalled in a state of chronic inflammation, and cannot progress through the later stages of healing (4–6). A hallmark characteristic of chronic inflammation is excess infiltration of neutrophils and macrophages. Leukocyte abundance can be harmful to wound healing due to high production of reactive oxygen species and serine proteases, which degrade structural proteins of the extracellular matrix therefore limiting proliferation and remodeling (7). In addition, diabetic wounds are colonized with numerous bacterial pathogens that trigger leukocyte influx and exacerbate poor healing outcomes (8).

Some of the most common species isolated from chronic wounds are *Staphylococcus aureus, Pseudomonas aeruginosa* and *Streptococcus agalactiae*, also known as Group B *Streptococcus* (GBS) (9–12). *S. aureus* is responsible for 76% of all skin and soft tissue infections and is found in wounds of both non-diabetic and diabetic individuals (11, 13). Thurlow et al. has shown that elevated glucose in hyperglycemic abscesses leads to increased virulence potential of *S. aureus*, and is dependent on two glucose transporters (14). In addition, specific *S. aureus* factors such as the proteases ClpX, ClpP and ClpC were shown to contribute to *S. aureus* skin infection in diabetic mice (15). *P. aeruginosa* is also a commonly isolated bacterium from wounds of both non-diabetic and diabetic individuals (11, 16). *P. aeruginosa* infection of chronic wounds has been linked to enhanced biofilm formation and activation of matrix metalloproteases that degrade extracellular matrix molecules (17, 18). Conversely, GBS is rarely identified from non-diabetic wounds, but is being increasingly identified in diabetic individuals (19, 20). While some work has characterized *S. aureus* and *P. aeruginosa* in diabetic wound infections, no previous studies have investigated GBS pathogenesis in the diabetic wound or on the skin in general. In addition, little is known about the mechanisms by which bacteria survive inflammation in the diabetic wound environment.

GBS is an opportunistic pathogen that is a leading cause of neonatal invasive infections. Approximately 20-30% of pregnant women are colonized with GBS in the vaginal tract, and can transmit this bacterium to the neonate *in utero* or during birth (21, 22). Neonatal infections can lead to the development of invasive diseases like sepsis, pneumonia and meningitis, with a mortality rate of 10-15% (23). In addition to neonatal infections, GBS infection of non-pregnant adults, including the immunocompromised is on the rise, with over two-thirds of invasive GBS disease occurring in adults (24). Of these individuals, the most common comorbidity is diabetes, which is present in 20-25% of nonpregnant adults with GBS (24). Despite this, the majority of GBS studies have focused on vaginal colonization and neonatal disease; and the mechanisms of GBS pathogenesis in diabetic infection have not been examined.

Numerous reports have identified mechanisms of GBS colonization that contribute to vaginal persistence, as well as mechanisms of invasive infection such as systemic disease and meningitis. GBS produces multiple virulence factors, including the β-hemolysin/carotenoid pigment (encoded by *cylE*), various surface adhesins such as pili, serine-rich repeat (Srr) proteins and BspC that bind host extracellular matrix proteins (ECMs) and receptors to promote adherence and invasion, as well as immune evasion factors such as the polysaccharide capsule, that allows the bacterium to resist phagocytic killing and persist in the human host (25–29). Ten capsular serotypes have been identified, with serotypes Ia, Ib, II, III and V being the most common serotypes associated with disease (30). The transition from a commensal lifestyle to infection is controlled by multiple regulators such as transcription factors and two-component systems (TCS) that coordinate the expression of these virulence factors in the host. TCS typically encode a sensor histidine kinase which recognizes a stimulus and a cognate response regulator which can be activated to control gene expression (31). GBS encodes approximately 21 unique TCS, which is more than other Streptococcal species such as *Streptococcus pneumoniae* and *Streptococcus pyogenes*, suggesting the importance of these regulators in this opportunistic pathogen (32, 33). Perhaps the most well-studies TCS in GBS is CovRS, encoding a sensor histidine kinase (*covS*) and a response regulator (*covR*). CovR represses the expression of multiple GBS virulence factors including fibrinogen and plasminogen binding proteins and the β-hemolysin (34). CovR regulation has been shown to be important for GBS vaginal colonization and development of invasive disease (35, 36).

Here, we describe for the first time the development of a murine model of GBS diabetic wound infection to begin to assess host-pathogen interactions in this environment. We find that GBS infection in the diabetic wound promotes a hyperinflammatory response with increased cytokine production and neutrophil degranulation. GBS responds to the inflammatory environment with the upregulation of numerous virulence factors such as *pbsP* and *cylE,* which are necessary for full GBS virulence in diabetic infection and acquiring mutations in *covR* during diabetic wound infection. Taken together, these results present numerous strategies by which GBS adapts to the hyperinflammatory diabetic wound environment and provides insight into the pathogenic mechanisms of GBS diabetic wound infection.

## Results

### Development of a Murine Model of GBS Diabetic Wound Infection

We developed a murine model of GBS diabetic wound infection using *Lepr^db^* mice. These mice harbor a mutation in the leptin receptor gene which results in obesity and mimics type 2 diabetes (37, 38). Mice were wounded with a 6 mm biopsy punch on their backs, infected with 1 x 10^7^ CFU of GBS and wrapped with an adhesive to allow efficient bacterial inoculation. Once the adhesive was removed, and animals were left for additional time before sacrifice, tissue collection and bacterial enumeration (**Fig 1A**). All mice exhibited non-fasting glucose levels above 250 mg/dL to be considered diabetic and levels under 200 mg/dL for non-diabetic controls (**Fig 1B**). Following GBS infection, diabetic animals had significantly larger wounds at the time of sacrifice than wild-type (WT) C57Bl/6J controls with the average wound size of non-diabetic mice being .097 cm^2^ and diabetic being .403 cm^2^ (p-value= .<.0001, standard deviations .047 and .177 respectively) (**Fig 1C,D**). In addition, we utilized three clinically relevant GBS strains representing the three of the five most prominent serotypes associated with disease worldwide; A909 (serotype Ia), COH1 (serotype III) and CJB111 (serotype V) (30). Regardless of strain, we found that even at an early time point of infection (one day after adhesive removal, four days after initial infection) there were significantly more bacteria recovered from the wounds of diabetic animals in comparison to the non-diabetic (33.9-fold for A909, 21.6-fold for COH1, and 2.6-fold for CJB111) (**Fig 1E**). We also performed experiments on both male and female mice and saw no significant differences in CFU recovered between sexes (**Supplemental Fig 1**), thus all subsequent infections were performed in female mice. In addition to GBS, we have recovered colonies of *Enterococcus faecalis* and *Staphylococcus xylosus* from wound homogenates. These species were only found in select mice and at low levels (> 10^2^ CFU/g wound). These data support clinical findings that GBS establishes infection in the diabetic wound environment (19, 39), and provide a relevant model for further investigation.

**Figure 1.**
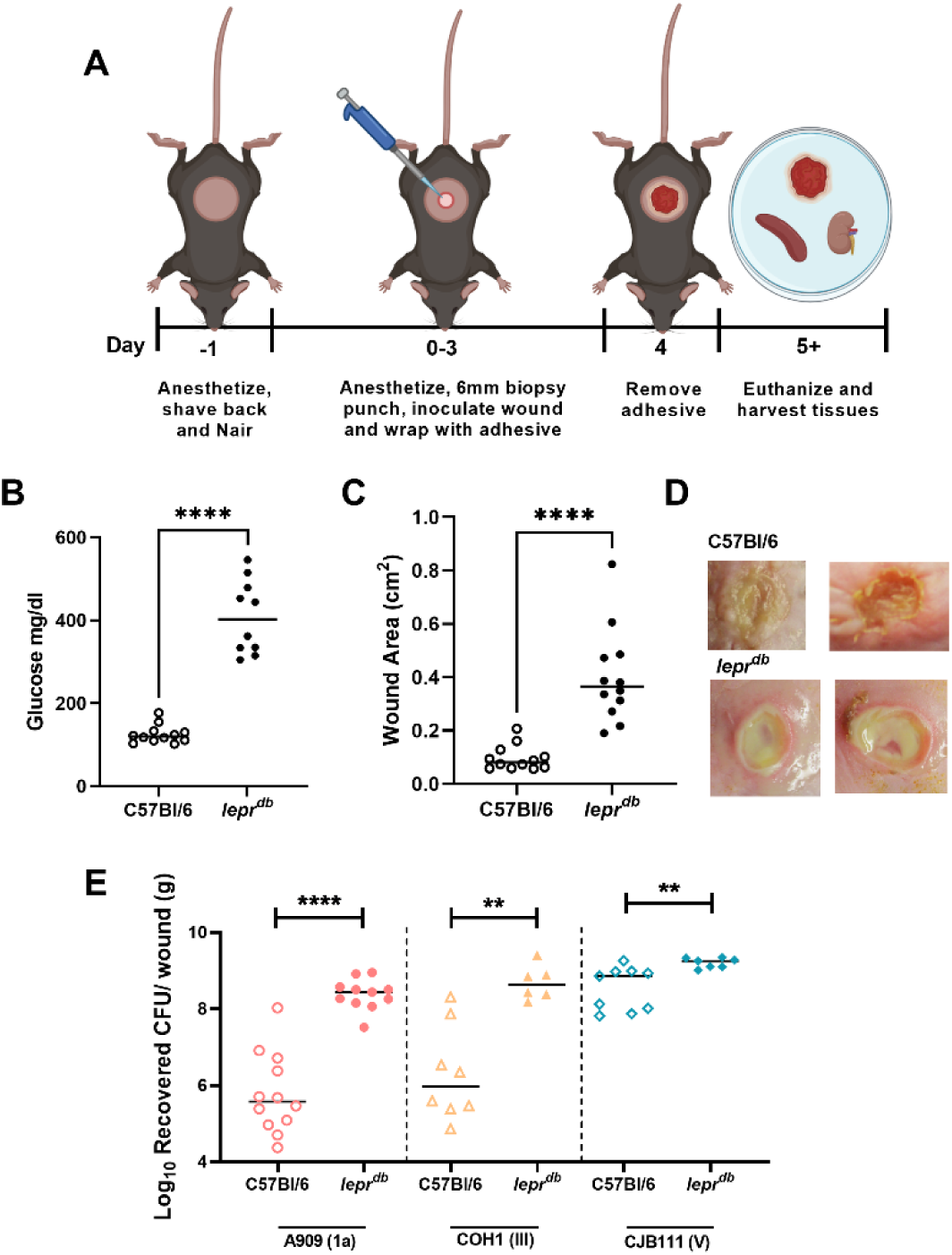
Murine Model of GBS Diabetic Wound Infection. (A) Schematic of the murine model of diabetic wound infection. (B) Non-fasting glucose levels from the blood of mice the day of infection. (C) Wound area calculated in ImageJ from representative mice following GBS infection. (D) Representative images of GBS infected wounds from non-diabetic and diabetic mice on the day of sacrifice. (E) CFU recovered from wounds of non-diabetic and diabetic mice after GBS infection. All animal infections proceeded for four days with three days under adhesive and sacrifice 24 h after adhesive removal. Significance determined by Mann–Whitney U test; *p<.05, **p<.01, ****p<.0001.

Further, we obtained 27 clinical isolates of GBS recovered from diabetic wounds of human patients, and determined the molecular serotype of each isolate using primers specific to *cps* loci (40). Serotypes Ia, Ib, II and III were confirmed by flow cytometry as described by Burcham et al. using monoclonal antibodies against capsule (41). The most prominent serotype was Ia, which represented one third of all isolates recovered (33.33%). The remaining isolates were serotype V (22.22%), II (22.22%), III (11.11%) and Ib (7.41%) with one isolate that was unable to be typed by either PCR or flow cytometry (**Supplemental Table 1**).

### The hyperinflammatory wound environment in diabetic mice is more susceptible to GBS persistence

To determine the host response during GBS infection, we performed dual RNA-sequencing of both diabetic and non-diabetic infected animals in comparison to animals inoculated with a PBS control. Principal component analysis (PCA) plots demonstrate that diabetic and non-diabetic animals clustered more similarly with infected vs. uninfected animals also clustering closely (**Fig 2A**). We utilized Reactome pathway analysis as well as gene set enrichment analysis (GSEA) to link upregulated genes to known cellular pathways (42). Transcriptomics performed on uninfected wounds support the current literature that *lepr^db^* mice promote upregulation of genes and pathways involving inflammation such as the interferon alpha response, interferon gamma response and inflammatory response in comparison to C57Bl/6J controls (**Supplemental Fig 2**) (43, 44).

**Figure 2.**
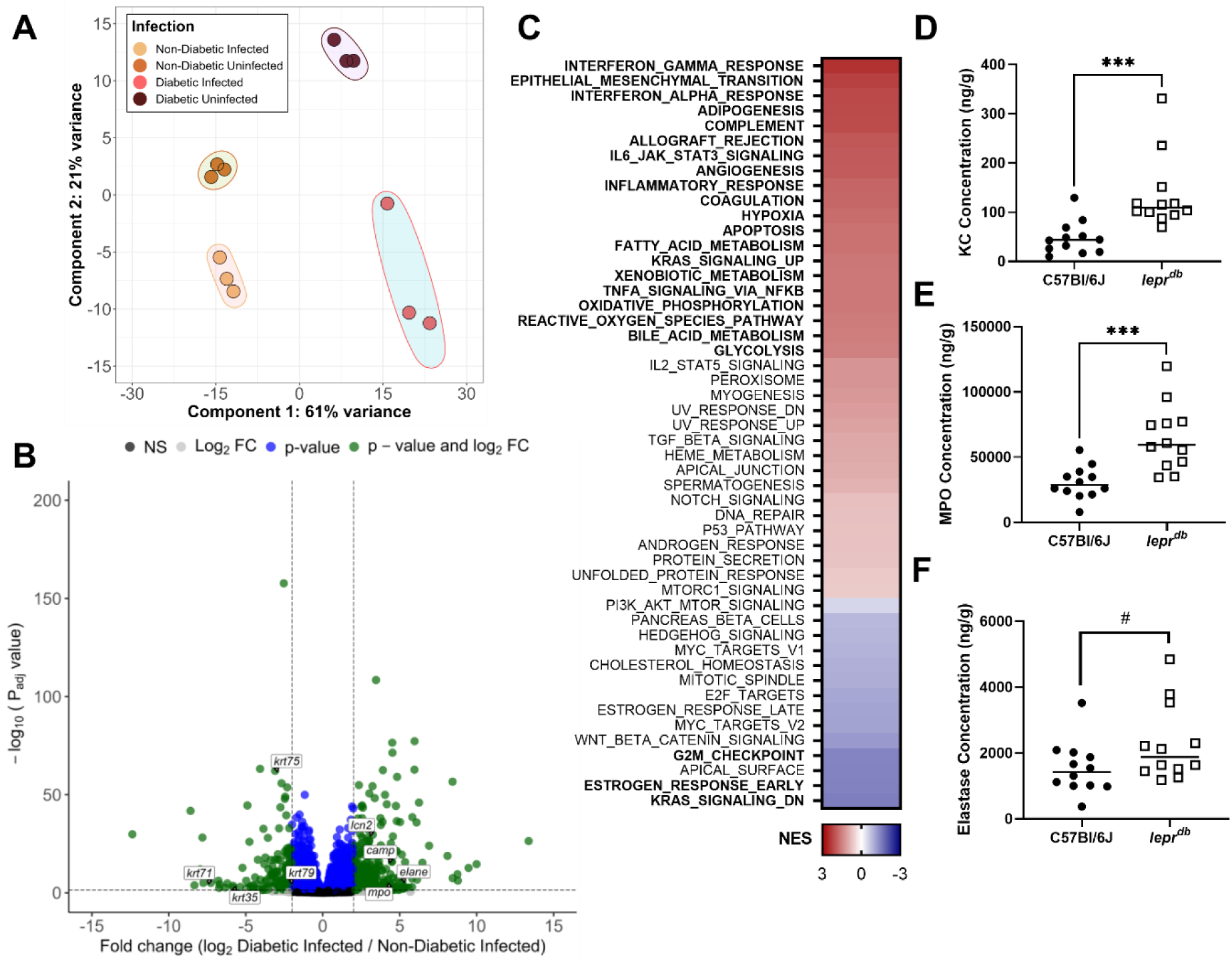
The hyper-inflammatory wound environment in diabetic mice is more susceptible to GBS persistence. (A) PCA plot of murine transcriptome from RNA-seq analysis. (B) Volcano plot of differentially expressed genes. (C) GSEA of pathways enriched in diabetic wounds infected with GBS in comparison to non-diabetic. Significant pathways are in bold. Normalized enrichment score (NES) presented as a heat map with highly enriched pathways in red. (D-F) ELISAs on tissue homogenates at time of sacrifice. All concentrations were normalized to tissue weight. Mice were infected for four days with three days under adhesive and sacrifice 24 h after adhesive removal. Significance determined by Mann-Whitney U-test; ***p<.001, # = .0597.

We next examined the transcriptomes of diabetic and non-diabetic mice infected with GBS to determine if the host response was altered in diabetic infection. A total of 326 transcripts were significantly upregulated, and 275 were significantly downregulated in diabetic infection in comparison to non-diabetic (Log2 fold-change > 2, FDR adjusted p-value< .05). Diabetic animals infected with GBS significantly upregulated genes involved in neutrophil degranulation, activation of matrix metalloproteases and antimicrobial peptides while downregulating genes involved in keratinization (**Fig 2B**, **Supplemental Table 2**).Pathway analysis of these genes revealed that GBS infection of diabetic wounds leads to the significant upregulation of numerous pathways including the interferon gamma response, the inflammatory response, and the reactive oxygen species pathway (**Fig 2C**). Enzyme-linked immunosorbent assays (ELISAs) were then performed on tissue homogenates from diabetic and non-diabetic animals infected with GBS. Wound tissues from diabetic mice had significantly higher abundance of the neutrophil chemoattractant KC (homologous to human CXCL1) and myeloperoxidase (MPO) and a marked increase in elastase (ELANE) in comparison to non-diabetic (**Fig 2D-F**).

### Diabetic animals upregulate inflammatory pathways and immune cell recruitment during GBS infection

After confirming that GBS survives better in the diabetic wound, we next sought to determine how infection with GBS affected gene expression specifically in diabetic mice. When comparing the transcriptome of diabetic animals infected with GBS compared to PBS controls, 146 transcripts were significantly upregulated, and 63 were significantly downregulated (Log2 fold-change > 2, FDR adjusted p-value< .05) with some of the most highly upregulated transcripts including the subunits of calprotectin (s1008a and s100a9), myeloperoxidase (*mpo),* the cathelicidin antimicrobial peptide (*camp*) and elastase (*elane*) (**Fig 3A**). Significantly upregulated pathways in GSEA included interferon gamma response, TNF alpha signaling via NFkB, complement, inflammatory response and reactive oxygen species (**Fig 3B**). NFkB activation has been linked to an amplified inflammatory state in chronic wounds due to increased production of inflammatory cytokines (7). We then ran our significantly dysregulated genes through Reactome and found some of the most highly upregulated pathways were neutrophil degranulation, IL-10 signaling and the formation of a fibrin clot (**Supplemental Table 3**). Interestingly, there were no pathways significantly downregulated in GSEA analysis, however, striated muscle contraction and myogenesis were both enriched in Reactome analysis (**Fig 3B, Supplemental Table 3**).

**Figure 3.**
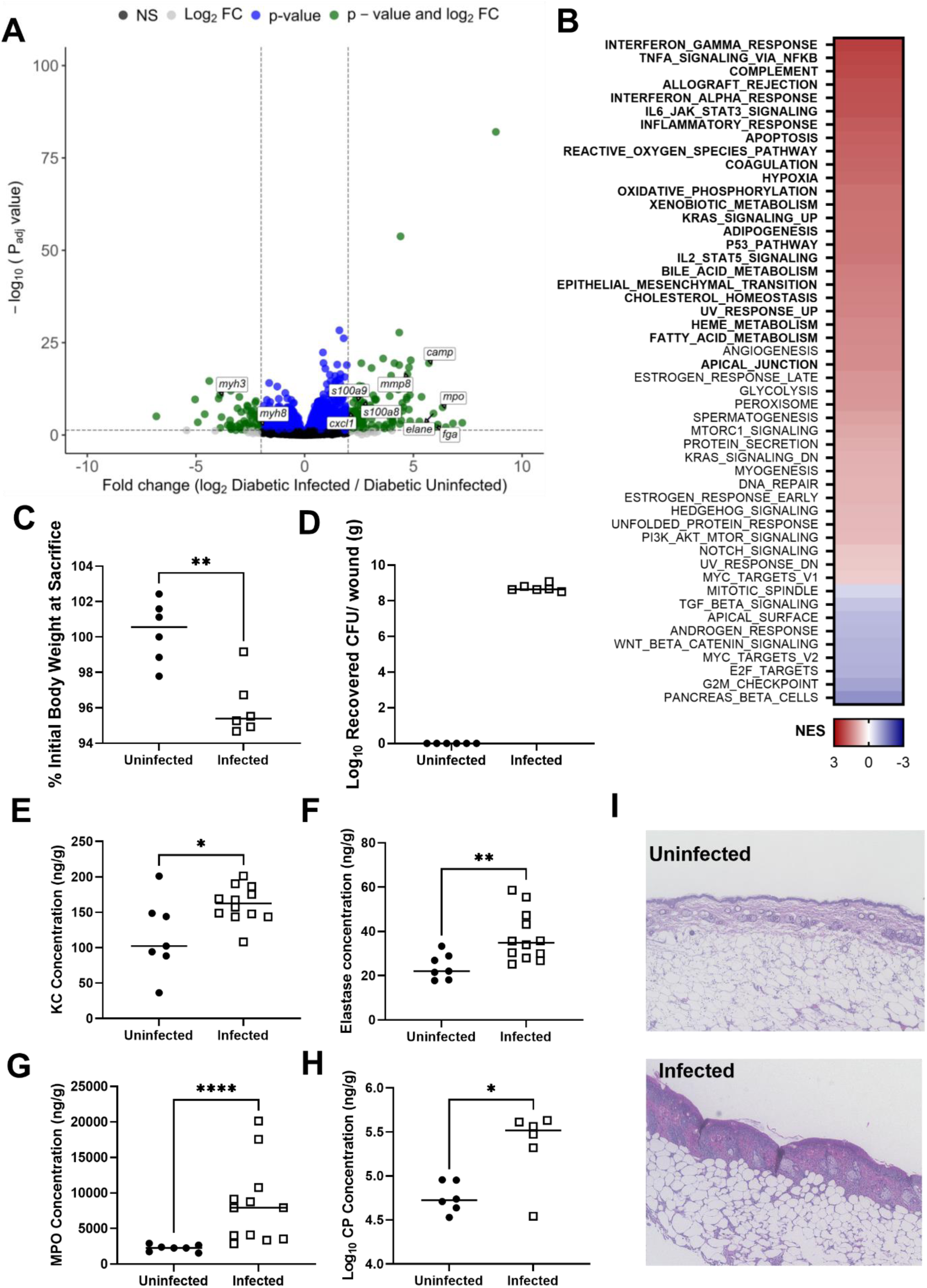
Diabetic animals upregulate inflammatory pathways and immune cell recruitment upon GBS infection. (A) Volcano plot for differentially expressed genes when comparing diabetic mice infected with GBS to uninfected controls. (B) Gene set enrichment analysis of diabetic animals infected with GBS vs. uninfected controls. Significant pathways in bold. NES presented as a heat map with highly enriched pathways in red. (C) Percent initial body weight of mice after GBS infection. (D) GBS recovery from wound tissues of uninfected vs. infected mice. (E-H) ELISAs on tissue homogenates at time of sacrifice. All concentrations were normalized to tissue weight. (I) Histology of diabetic wound tissue at time of sacrifice. Mice were infected for four days with three days under adhesive and sacrifice 24 h after adhesive removal. Significance determined by Mann-Whitney U-test; *p<.05, **p<.01, ****p<.0001.

The upregulation of inflammatory pathways led us to hypothesize that GBS triggers increased inflammation in the diabetic wound. Mice infected with GBS lost significantly more weight over the course of infection than uninfected controls, and none of the uninfected mice had any GBS recovered from wound tissue (**Fig 3C,D**). As mentioned above we used ELISA to determine that GBS infected wounds had significantly greater abundance of the neutrophil chemoattractant KC further supporting that GBS infection triggers the recruitment of neutrophils to the site of infection (**Fig 3E**). Finally, we determined that the abundance of neutrophil components elastase, myeloperoxidase and a subunit of calprotectin are significantly higher during GBS infection (**Fig 3F-H**). H&E staining of wound tissue sections demonstrate a gross difference in tissue architecture from GBS infected mice including thickening of the epithelial layer and inflammatory infiltrate (**Fig 3I**). Collectively, these data suggest that GBS promotes inflammation in the already highly inflammatory diabetic wound environment and confirms the upregulation of these pathways and genes observed in our RNA-seq analysis (**Fig 3A,B**, **Supplemental Table 3**).

### GBS transcriptome in the diabetic wound

To characterize the bacterial response in the diabetic wound environment, we performed RNA-sequencing on GBS isolated from non-diabetic and diabetic wounds in comparison to GBS grown *in vitro* (input control). PCA plots indicate that each group clusters independently, with the largest variation being between the input control bacteria and the bacteria isolated from the mice (**Fig 4A**). A total of 974 transcripts were significantly altered (minimum fold change of 3, FDR adjusted p-value <.05) when comparing GBS isolated from diabetic infection in comparison to the input control with 461 being upregulated and 513 downregulated (**Fig 4B**). Further analysis of genes with altered regulation during diabetic infection revealed multiple housekeeping genes were downregulated during GBS infection including ribosomal subunit genes *rpsC*, *rplV* and *rpsE*. (**Fig 4C**). In addition, multiple genes with predicted roles in lipid transport were downregulated in diabetic infection. Conversely, numerous virulence associated genes were upregulated in comparison to the input control (**Fig 4C**, **Table 1**). Some of the most highly upregulated transcripts in diabetic infection include the gene encoding the surface plasminogen binding protein PbsP, the *cyl* operon encoding GBS hemolysin and pigment, quorum sensing peptide pheromone *shp2*, nuclease and protease encoding genes such as *nucA* and *clpL*, as well as various predicted effectors of the type VII secretion system (**Table 1**.) Interestingly, many of these factors are known to be part of the core regulon of the CovRS TCS (45–47).

**Figure 4.**
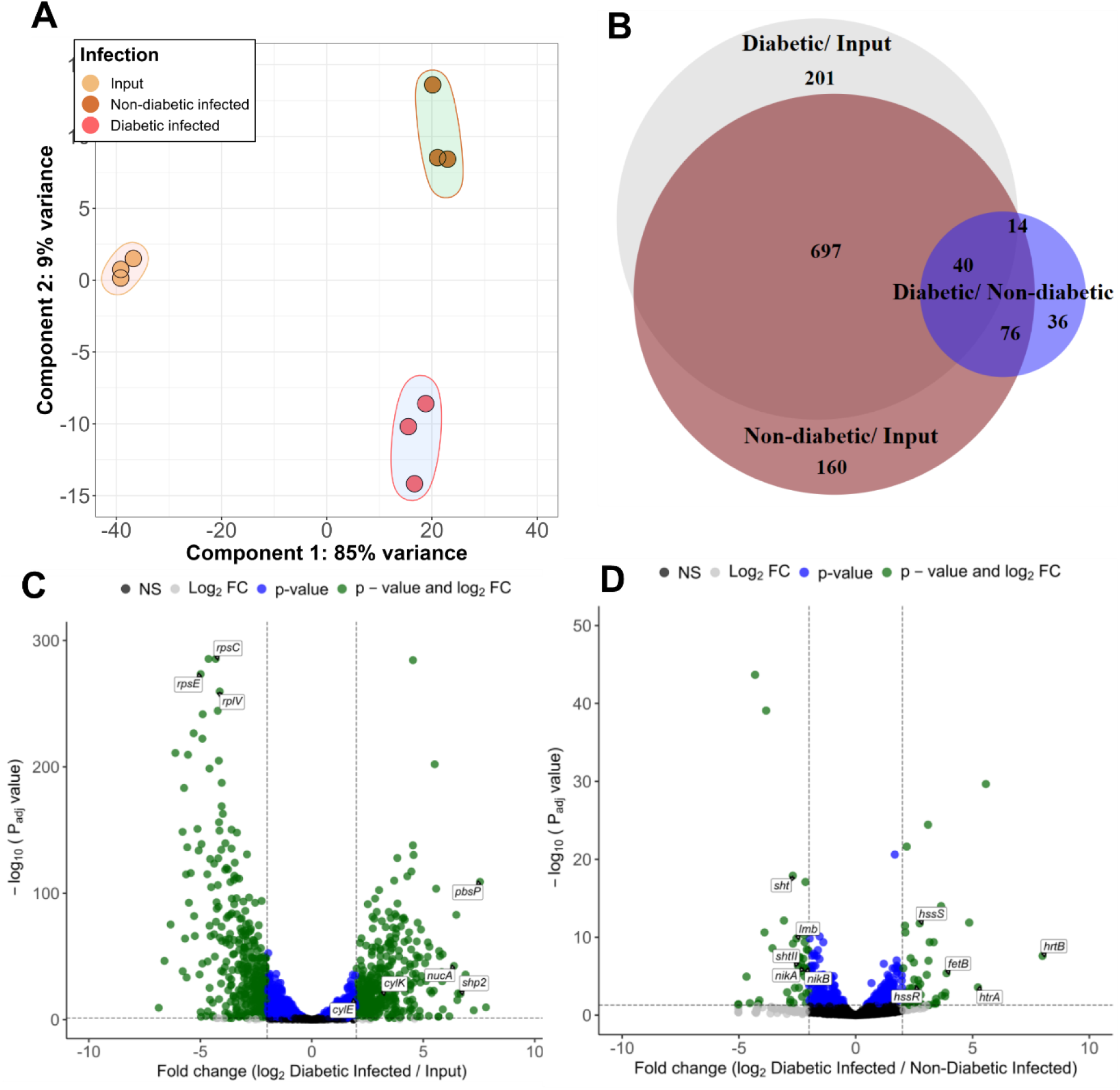
GBS Transcriptome in Diabetic Wound Infection. (A) PCA plot of bacterial transcriptome in RNA-seq. (B) Venn diagram of differentially expressed genes in each comparison. (C) Volcano plot of differentially expressed genes in diabetic infection vs. input control. (D) Volcano plot of differentially expressed genes in diabetic vs. non-diabetic wound infection.

**Table 1.**
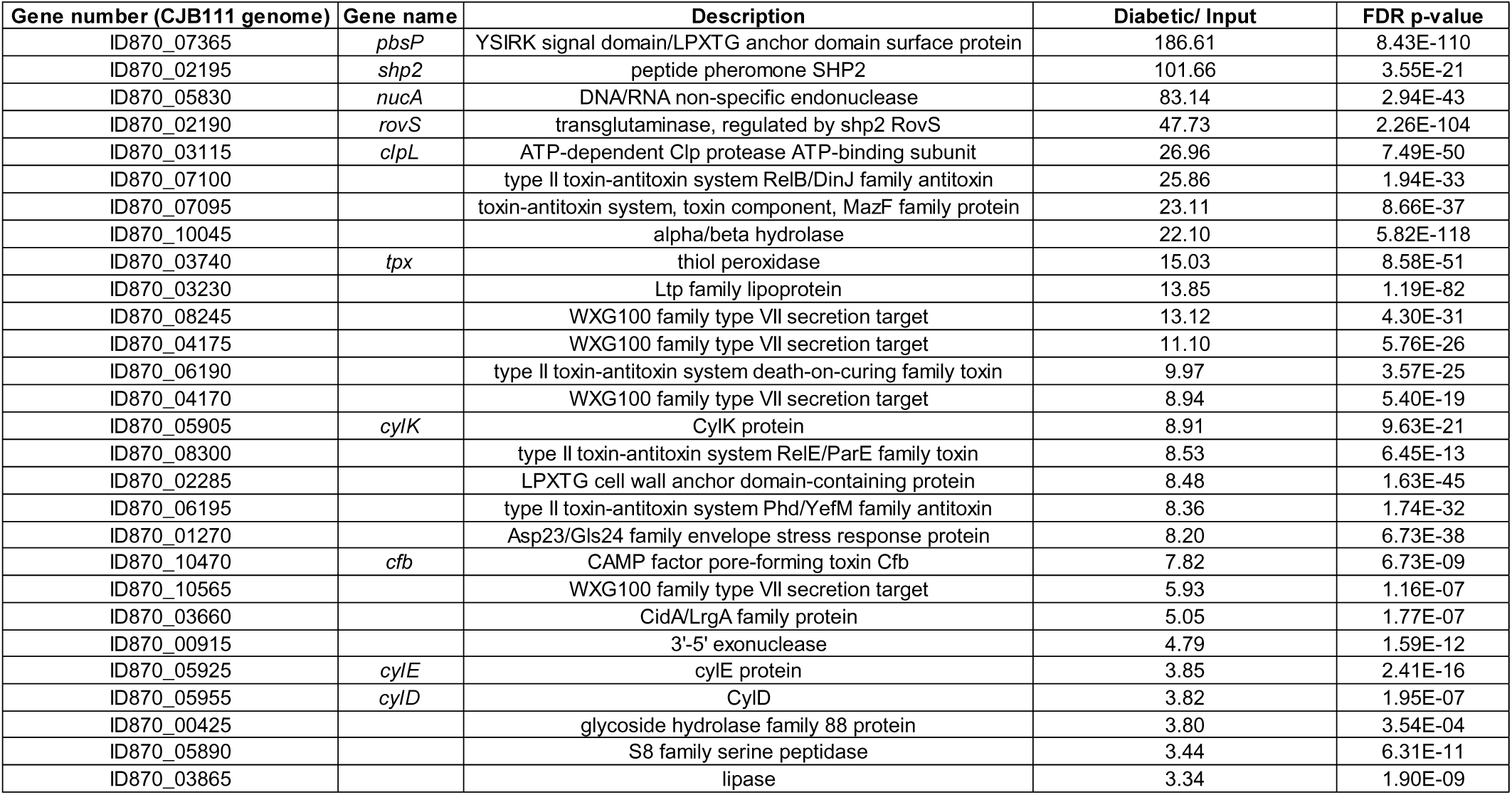
Select GBS Virulence factors upregulated in diabetic infection vs. input control.

| Gene number (CJB111 genome) | Gene name | Description | Diabetic/ Input | FDR p-value |
| --- | --- | --- | --- | --- |
| ID870_07365 | *pbsP* | YSIRK signal domain/LPXTG anchor domain surface protein | 186.61 | 8.43E-110 |
| ID870_02195 | *shp2* | peptide pheromone SHP2 | 101.66 | 3.55E-21 |
| ID870_05830 | *nucA* | DNA/RNA non-specific endonuclease | 83.14 | 2.94E-43 |
| ID870_02190 | *rovS* | transglutaminase, regulated by shp2 RovS | 47.73 | 2.26E-104 |
| ID870_03115 | *clpL* | ATP-dependent Clp protease ATP-binding subunit | 26.96 | 7.49E-50 |
| ID870_07100 |  | type II toxin-antitoxin system RelB/DinJ family antitoxin | 25.86 | 1.94E-33 |
| ID870_07095 |  | toxin-antitoxin system, toxin component, MazF family protein | 23.11 | 8.66E-37 |
| ID870_10045 |  | alpha/beta hydrolase | 22.10 | 5.82E-118 |
| ID870_03740 | *tpx* | thiol peroxidase | 15.03 | 8.58E-51 |
| ID870_03230 |  | Ltp family lipoprotein | 13.85 | 1.19E-82 |
| ID870_08245 |  | WXG100 family type VII secretion target | 13.12 | 4.30E-31 |
| ID870_04175 |  | WXG100 family type VII secretion target | 11.10 | 5.76E-26 |
| ID870_06190 |  | type II toxin-antitoxin system death-on-curing family toxin | 9.97 | 3.57E-25 |
| ID870_04170 |  | WXG100 family type VII secretion target | 8.94 | 5.40E-19 |
| ID870_05905 | *cylK* | CylK protein | 8.91 | 9.63E-21 |
| ID870_08300 |  | type II toxin-antitoxin system RelE/ParE family toxin | 8.53 | 6.45E-13 |
| ID870_02285 |  | LPXTG cell wall anchor domain-containing protein | 8.48 | 1.63E-45 |
| ID870_06195 |  | type II toxin-antitoxin system Phd/YefM family antitoxin | 8.36 | 1.74E-32 |
| ID870_01270 |  | Asp23/Gls24 family envelope stress response protein | 8.20 | 6.73E-38 |
| ID870_10470 | *cfb* | CAMP factor pore-forming toxin Cfb | 7.82 | 6.73E-09 |
| ID870_10565 |  | WXG100 family type VII secretion target | 5.93 | 1.16E-07 |
| ID870_03660 |  | CidA/LrgA family protein | 5.05 | 1.77E-07 |
| ID870_00915 |  | 3'-5' exonuclease | 4.79 | 1.59E-12 |
| ID870_05925 | *cylE* | cylE protein | 3.85 | 2.41E-16 |
| ID870_05955 | *cylD* | CylD | 3.82 | 1.95E-07 |
| ID870_00425 |  | glycoside hydrolase family 88 protein | 3.80 | 3.54E-04 |
| ID870_05890 |  | S8 family serine peptidase | 3.44 | 6.31E-11 |
| ID870_03865 |  | lipase | 3.34 | 1.90E-09 |

We next compared the bacterial transcriptome from GBS recovered from diabetic wounds vs. non-diabetic wounds. A total of 166 genes had significantly altered expression in this comparison, with 75 genes being upregulated and 91 being downregulated in diabetic infection in comparison to non-diabetic (**Fig 4B,D**). Multiple genes involved in sugar transport were downregulated in bacteria recovered from diabetic wounds, as well as genes involved in metal transport of iron, zinc, and manganese (**Supplemental Table 4**). Conversely, many genes involved in iron export were upregulated in bacteria recovered from diabetic wounds (**Supplemental Table 4**).

### The CovRS regulon contributes to diabetic wound infection

We observed when plating GBS recovered from the diabetic wound that many colonies were hyperpigmented (**Fig 5A**). These phenotypes were stable, and colonies were hyperhemolytic when plated on sheep’s blood agar (**Fig 5B**). Interestingly, we have only observed this phenotype in colonies recovered from diabetic wounds with ∼60% (7/12) of diabetic wound homogenates containing a hyperpigmented colony compared to 0% (0/12) in non-diabetic wounds. We next infected mice with GBS and sacrificed either 24 h or 1 week after adhesive removal and noticed a marked increase in the total number of pigmented colonies recovered over time (**Fig 5C**) suggesting that *covR* mutations are selected for in the diabetic wound, specifically. Hyperpigmentation and hyperhemolysis in numerous GBS strains has been attributed to mutations in the *covRS* TCS, due to the subsequent de-repression of the *cyl* operon which is linked to GBS hemolysis and pigment production (25, 48–52). We surveyed the *covR* locus from hyperpigmented/ hyperhemolytic strains recovered from 13 different mice for single nucleotide polymorphisms (SNPs) and observed multiple SNPs resulting in mutations in the *covR* locus. Of these, 100% of the colonies encoded *covR* mutations with 8/13 encoded for amino acid substitutions in the receiver domain of CovR, 3/13 containing single-nucleotide deletions and 2/13 encoding insertions or deletions (**Fig 5C, Supplemental Table 5**). Nonpigmented colonies were also isolated and sequenced from each mouse containing a hyperpigmented colony and none of the nonpigmented colonies had mutations in *covR*. We hypothesized due to the upregulation of the *cyl* operon in diabetic wound infection and the spontaneous *covR* mutations *in vivo*, that a *ΔcylE* mutant would be attenuated in diabetic wound infection. In confirmation, we infected diabetic mice with the *ΔcylE* mutant strain and observed a significant decrease in bacteria recovered from wound tissue compared to WT infected mice (**Fig 5D**). Further, wounds from *cylE* infected mice had a significant reduction in the abundance of the chemokine KC (CXCL1) (**Fig 5E**). As expected, we saw no significant differences in bacterial recovery when comparing a clean deletion *ΔcovR* mutant strain to our WT *in vivo* (**Supplemental Fig 3**), likely due to the fact that WT GBS is acquiring *covR* mutations in the wound.

**Figure 5.**
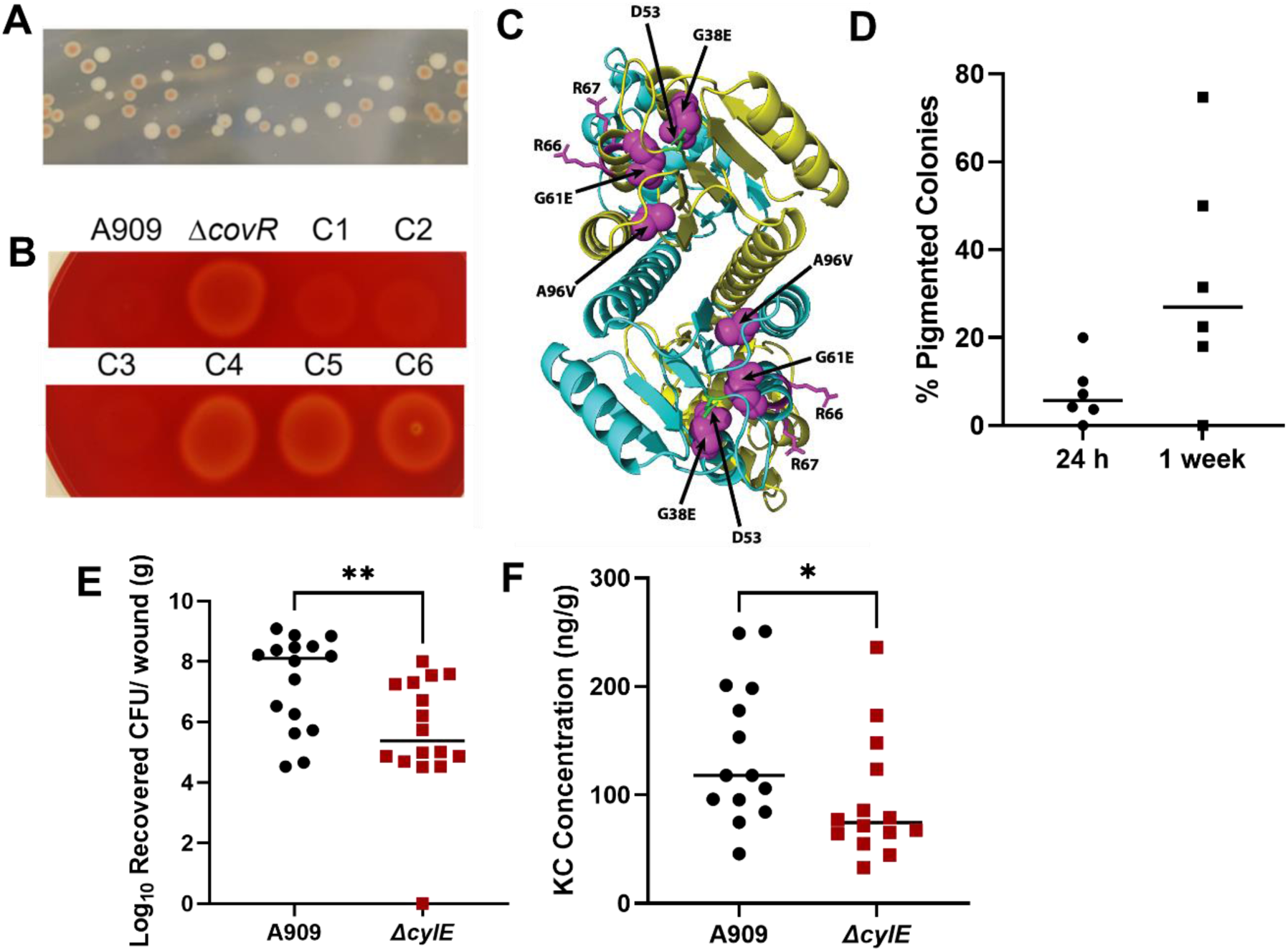
The CovRS Regulon Contributes to Diabetic Wound Infection. (A) GBS colonies on THA recovered from diabetic wounds after four days of infection. (B) Hyperpigmented colonies plated on sheep’s blood agar; C1-3 from non-pigmented colonies and C4-6 from pigmented colonies (C) Structural model of CovR mutations on VanR structure from *Streptomyces coelicolor.* (D) Percentage of total GBS colonies from diabetic wound tissue 24 h or 1 week after adhesive removal which were hyper-pigmented on Todd-Hewitt agar. (E) CFU of GBS recovered from diabetic wound tissue 24 h after adhesive removal. (F) KC concentration recovered from wound tissue homogenates. All ELISA data is normalized to tissue weight. Significance determined via Mann Whitney U-test; *p<.05, **p<.01.

### The surface protein PbsP contributes to diabetic wound formation via adherence to the skin

The most highly upregulated gene during GBS diabetic wound infection encodes the surface plasminogen binding protein PbsP (**Table 3**). Interestingly, *pbsP* has been shown to be repressed by CovR in multiple GBS strains (26, 34, 53). In addition, PbsP is known to bind multiple extracellular matrix components such as plasminogen and fibrinogen (26) and contribute to adherence to colon epithelial and brain endothelial cells (26, 54, 55) We therefore hypothesized that PbsP contributes to diabetic wound infection by promoting adherence to the skin. A Δ*pbsP* mutant is significantly attenuated in diabetic wound infection in comparison to WT (**Fig 8A**). Further, Δ*pbsP* mutant infected tissues have significantly less abundance of neutrophil markers MPO, elastase and calprotectin, which are all upregulated during GBS diabetic wound infection (**Table 2**) (**Fig 8B-D**). We next tested the ability of the Δ*pbsP* mutant to adhere to skin cells *in vitro*. Using a cell culture line of immortalized keratinocytes (NTERTs), we found that the Δ*pbsP* mutant had a ∼50% reduction in adherence to skin epithelial cells that could be restored by complementing the mutant with WT *pbsP* (**Fig. 8E**) (56). Additionally, we tested whether PbsP or CylE contributes to GBS infection of non-diabetic wounds and found no significant difference in bacterial recovery (**Supplemental Fig 4**). These data suggest that PbsP contributes to diabetic wound infection and direct interaction with skin keratinocytes.

## Discussion

A hallmark characteristic of diabetic individuals is an altered neutrophil response due to persistent hyperglycemia. Neutrophils from diabetic individuals are often pro-inflammatory, exhibiting increased neutrophil extracellular trap (NET) formation, pro-inflammatory cytokine production and extracellular ROS generation (1, 57). Additionally, decreased apoptosis, neutrophil migration and intracellular ROS production have been shown in diabetic individuals, leading to impaired bacterial killing (58). Persistent inflammation and decreased bacterial clearance are detrimental to diabetic individuals who develop wounds, as resolution of inflammation is paramount to proper wound healing (59). GBS is one of the most commonly isolated pathogens from diabetic wounds, and has been shown to promote inflammation and neutrophil influx during lung infection as well as during sepsis (11, 50, 60). Here, we have utilized dual RNA-sequencing to determine the transcriptional consequences of GBS infection of non-diabetic and diabetic wounds on both the host and pathogen. We demonstrate that GBS exacerbates inflammation in the already hyperinflammatory diabetic wound environment and identify the first mechanisms by which GBS adapts to this hyperinflammation (**Fig. 7**). The data herein are the first to provide insight into GBS pathogenesis in this clinically relevant model of infection.

**Figure 6.**
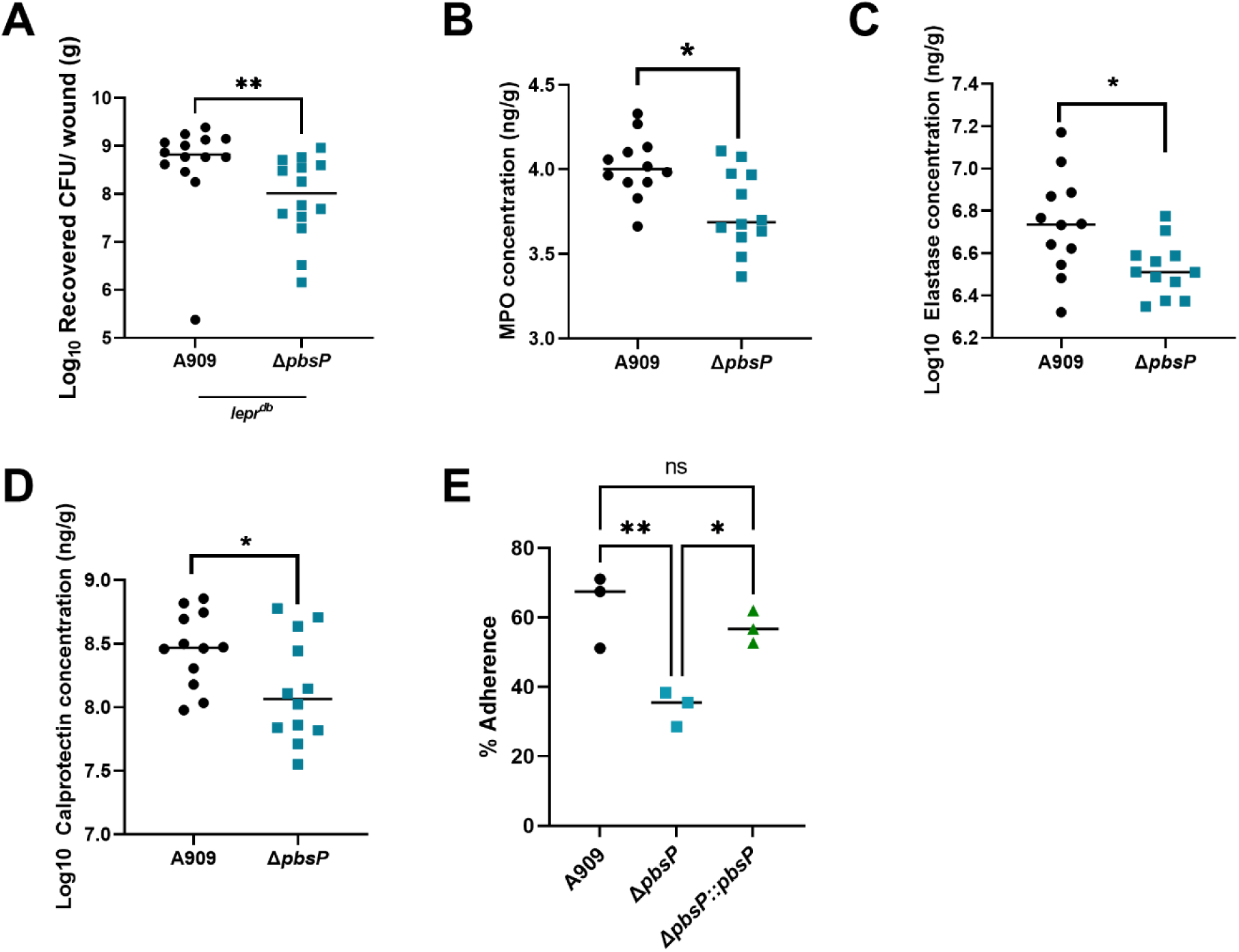
PbsP contributes to GBS burden and inflammation in the diabetic wound. (A) CFU recovered from diabetic mice infected with A909 or a *ΔpbsP* mutant after four days of infection. (B,C,D) MPO, Elastase, and Calprotectin concentration recovered from wound tissue homogenates. All ELISA data is normalized to tissue weight. (E) GBS adherence to immortalized keratinocytes with either wildtype, mutant or complemented PbsP. Significance determined via Mann Whitney U-test; *p<.05, **p<.01.

**Figure 7.**
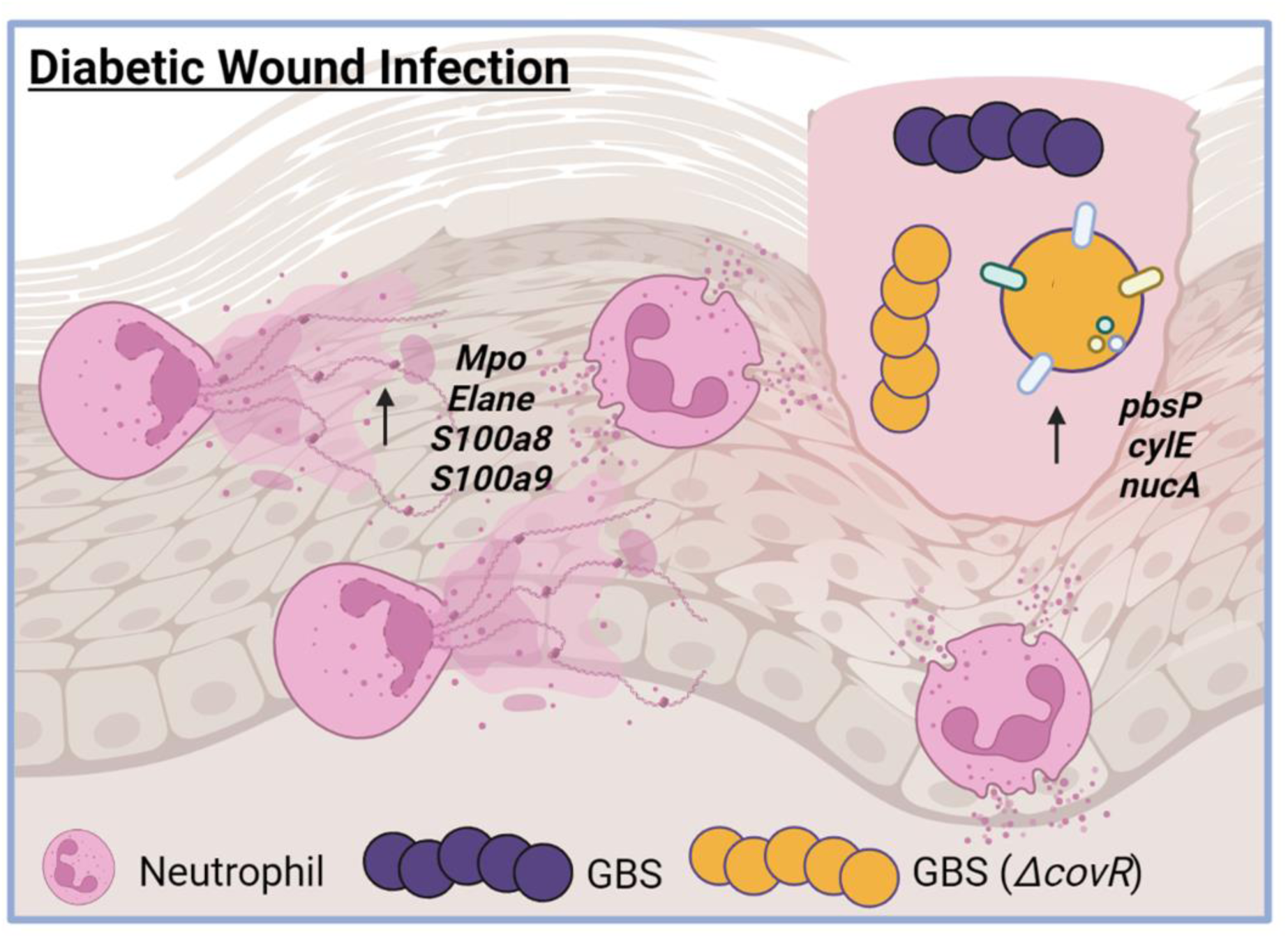
Model for GBS pathogenesis in the diabetic wound. Image generated in BioRender.

We demonstrate that GBS burden is significantly greater in diabetic wounds than non-diabetic, and that these data remain consistent regardless of GBS capsular serotype. GBS infection of diabetic wounds resulted in significant upregulation of pathways involved in the inflammatory response and ROS production in comparison to diabetic wounds without GBS. These findings are particularly interesting as diabetic animals already exhibit significant upregulation of these pathways in comparison to non-diabetic controls, suggesting that the presence of GBS exacerbates inflammation, particularly neutrophil recruitment and activation, during diabetes. We speculate that GBS promotes inflammation as a strategy to stall wound healing and closure. The presence of microbes has been linked to neutrophil influx in chronic wounds, but studies on specific bacterial factors that assist in promoting and surviving inflammation are limited.

An additional hypothesis is that the altered neutrophil response in diabetic individuals is a large reason GBS can survive the diabetic wound environment. Work by Thurlow et al. demonstrated that diabetic mice infected with *S. aureus* had impaired phagocyte function including decreased respiratory bursts and increased glucose available for the bacterium (14). While the contribution of glucose to GBS pathogenesis is not completely understood, it is known that increased glucose leads to thickening of the GBS capsule (61), which may contribute to virulence or survival in the wound. It is also possible that additional changes in neutrophil function change host nutritional immunity with respect to metal availability and sequestration. While we see increased calprotectin in diabetic wounds infected with GBS, we do not know if it is functional in metal sequestration. Future studies on metal availability in the diabetic wound will be critical in understanding how host nutritional immunity may impact GBS persistence.

Our bacterial RNA-sequencing revealed that GBS upregulates numerous virulence factors during diabetic wound infection in comparison to those grown in laboratory media. The most highly upregulated GBS gene in diabetic wound infection encodes for the surface plasminogen binding protein PbsP, which binds to extracellular matrix components such as plasminogen and fibrinogen (26). Plasminogen binding proteins are known to bind plasminogen, and, in the presence of a tissue activator, activate plasminogen into plasmin (62). While plasminogen is crucial in early wound healing to promote inflammation, it’s resolution is necessary to progress tissue healing into the proliferation phase (63, 64). A Δ*pbsP* mutant is attenuated in multiple GBS models of infection including in vaginal colonization and meningitis (54, 65). Furthermore, Lentini et al. demonstrated that brain tissue homogenates from mice infected with a Δ*pbsP* mutant had a significant reduction in MPO, TNFα and IL-1β abundance in comparison to WT infected mice (54). Here, we demonstrate that PbsP is required for GBS diabetic wound infection, and that Δ*pbsP* mutant infected wounds have significantly less MPO, elastase and calprotectin than those infected with WT. We hypothesize that PbsP binding to host plasminogen may be necessary for GBS mediated inflammation via increased cytokine and chemokine signaling as well as increased plasmin production that degrades host matrix components necessary for proliferation and remodeling of the skin. Further work on the exact mechanism of PbsP mediated inflammation is being elucidated.

We also investigated the GBS hemolysin/ pigment encoded by the *cyl* operon, which was highly upregulated during diabetic infection and is known to be part of the CovRS regulon (50, 51). Studies on the *cylE* component of this operon have shown that CylE contributes to GBS pathogenesis in murine models of sepsis and lung infection, and that induction of the neutrophil chemoattractant IL-8 was significantly reduced in A549 cells infected with a *cylE* mutant (50, 51). Diabetic wounds from mice infected with a *ΔcylE* mutant had significantly fewer bacteria recovered than those infected with WT GBS, coupled with significantly less abundance of the neutrophil chemoattractant KC (CXCL1). CXCL1 is known to attract neutrophils to the site of infection, and the abundance of CXCL1 in chronic wounds is high in comparison to burn or surgical wounds (66). We therefore hypothesize that production of the GBS hemolysin/pigment is necessary for GBS to trigger neutrophil influx into the diabetic wound environment.

Other factors that were highly upregulated during diabetic wound infection included the nuclease *nucA*, and the quorum sensing (QS) genes *shp2* and *rovS*. NucA is the major GBS nuclease and has been shown to degrade NETs as well as contribute to virulence in the lung (67). We speculate that the upregulation of *nucA* might be in response to the hyperinflammatory environment, and in particular, the increased NET production which is associated with diabetic wounds (1, 57). Increased nuclease activity could therefore assist GBS in evading NETs and persisting in the diabetic wound environment, although further experiments are warranted to determine this. Increased production of *shp2* and *rovS* suggests a need for bacterial cell-to-cell communication in the diabetic wound environment. The need for QS systems in wounds is well-documented in both *S. aureus* and *P. aeruginosa*, which both rely on QS systems for full virulence and persistence in chronic wounds (68, 69). While GBS *shp2* and *rovS* are required for GBS to persist in the murine liver and spleen (70), no work has investigated whether this QS system is critical to survival in the wound environment. Additionally, GBS *shp2* and *rovS* have a known role in interspecies communication which could highlight the importance of these genes in a polymicrobial diabetic wound environment (71).

When comparing the GBS transcriptome in diabetic vs. non-diabetic infection, we noticed a striking number of genes involving transport were dysregulated. Some of the most highly upregulated genes included *htrAB*, encoding the heme-regulated efflux pump and *hssRS*, the two-component system that regulates *hrtAB* expression. These systems are critical for GBS survival as they prevent a lethal heme overdose (72). Frank et al. demonstrated that heme accumulates at the wound edge of rats in early infection, and that this leads to recruitment of leukocytes to infection (73). We speculate, since the diabetic wound is hyperinflammatory, that there is excess heme in this environment, and GBS upregulates heme efflux systems to survive potential toxicity. Other downregulated genes include components of iron, zinc, and sugar transport systems. We hypothesize that GBS is turning these transport genes off, as there may be excess metals and sugar in the diabetic wound environment. Thus, it is possible that the diabetic host may not be successfully sequestering metals in the wound thereby allowing for GBS persistence, but this warrants further investigation.

Finally, we noticed that GBS colonies recovered from diabetic wounds were often hyperpigmented and encoded mutations in the TCS *covRS*. CovR represses numerous GBS virulence factors including the *cyl* operon as well as *pbsP* in multiple GBS strains (25, 35, 46). Of note, is that spontaneous mutations in the *covRS* locus of GBS have been identified in clinical isolates from patients with necrotizing fasciitis, sore throat, and prosthetic joint infection (48, 49, 74). In addition, numerous studies of *Streptococcus pyogenes* (GAS) have linked spontaneous mutations in *covRS* to invasive infection in this pathogen (75, 76). Importantly, we have never recovered any hyperpigmented colonies from non-diabetic animals. We therefore speculate that the diabetic environment is promoting the selection of *covR* mutants in the wound. The exact environmental factors that may promote *covR* mutations in the diabetic wound are currently unknown. Interestingly, Jorge et al. found that the antimicrobial peptide LL-37 can bind to CovS in GAS, resulting in the up-regulation of virulence factors (77). It is possible that GBS CovS may also bind to antimicrobial peptides such as those we have shown are upregulated in the diabetic wound environment. If this is the case, increased antimicrobial peptides in the diabetic environment may be driving GBS to increase virulence factor production, and contribute to the selection of *covR* mutants in the diabetic wound. It will also be of interest in future studies to examine the possibility that mutations may also occur in *covS*, which could differentially impact GBS persistence in the diabetic wound environment.

Using our model for GBS diabetic wound infection, we define changes in both the host and pathogen transcriptome in the murine diabetic wound environment and demonstrate that GBS promotes inflammation which contributes to the chronic wound environment. As we demonstrate, this model is an efficient tool to investigate GBS pathogenesis, and potentially other pathogens, in the context of diabetic wounds. Our results emphasize the importance of further studies on GBS pathogenesis and the innate immune response at the diabetic wound interface.

## Materials and Methods

### Bacterial strains and growth conditions

GBS strains A909, COH1 and CJB111 were used in this study. GBS strains were grown in Todd Hewitt Broth (THB; Research Products International, RPI) statically at 37° C. When needed, antibiotic was added to THB at final concentrations of 100 μg/mL spectinomycin. Strains containing the plasmid pDCErm were grown in THB + 5 μg/mL erythromycin. All strains used in this study can be found in **Supplemental Table 6** and primers in **Supplemental Table 7.** The human immortalized keratinocyte cell line N/TERT-2G cells were kindly gifted from Dr. Johann Gudjonsson with permission from Dr. Jim Rheinwald and were maintained up to 25 passages at 37 ° C with 5% CO_2_ according to methods described in Dickson et al., 2000 (56).

### Ethics statement

Animal experiments were approved by the Institutional Animal Care and Use Committee (IACUC) at University of Colorado Anschutz Medical Campus protocol #00987 and were per-formed using accepted veterinary standards. The University of Colorado Anschutz Medical Campus is AAALAC accredited; and its facilities meet and adhere to the standards in the “Guide for the Care and Use of Laboratory Animals”.

### Murine model of wound infection

Female or male, 8-week-old C57Bl/6J mice or *Lepr^db^* mice were anesthetized with isoflurane the day prior to infection. The backs of the mice were shaved and treated with Nair and tail snips were performed to measure blood glucose levels with a glucometer. The next day mice were weighed and again anesthetized before undergoing wounding procedure. The backs of mice were disinfected with iodine wipes and then wounded with a 6 mm biopsy punch. Following, 1 x 10^7^ CFU of GBS or PBS control was added before wrapping the mice in the adhesive Tegaderm^TM^. Wounds were left for 72 h before removal of the adhesive. After another 24 h, mice were sacrificed via CO_2_ inhalation and tissues were removed and homogenized for bacterial enumeration. Tissues were placed 500 µL of PBS in 2.0 mL conical screw cap tubes (Fisher) with 1.0 mm diameter zirconia/silica beads [BioSpec Cat. No. 1107911] and homogenized by bead beating two times for 60 sec in a BioSpec mini bead beater. Tissue homogenates were plated on GBS CHROMagar [SB282(B)], which allows only for the growth of GBS (in pink) and *Enterococcus* spp. (in blue). These experiments were approved by the committee on the use and care of animals at the University of Colorado-Anschutz Medical Campus in our protocol #00987.

### Histology

Mice were wounded as described above. Following sacrifice, half of the wound tissue was removed and fixed in formalin for 48 h. After, formalin was removed and wounds were placed into 70% EtOH and sent to the histology core for processing at the Gates Center for Regenerative Medicine, Dermatology at CU Anschutz. Images were taken with a BZ-X710 microscope (Keyence).

### RNA preparation and RNA-sequencing

Mice were wounded as previously described and tissues were collected and placed into 500 µL of RNA protect from the Qiagen RNeasy kit in 2.0 mL conical screw cap tubes with 1.0 mL zirconia beads as described above. Samples were placed into a bead beater and homogenized three times for 60 sec with five minutes of ice between each bead beating step. Sample supernatants were then collected and placed into new tubes for centrifugation. Samples were centrifuged for 10 minutes at 13,000 RPM and the pellet was resuspended in 600 µL of RLT plus β-Mercaptoethanol and homogenized by beat beating for 60 sec with .1 mL zirconia beads. Resulting homogenates were used for RNA preparation with the Qiagen RNeasy kit. For mouse samples, RNA was extracted from 2-3 mice before pooling RNA to form triplicate samples. For bacteria, we utilized three input controls as replicates and again pooled from 2-3 mice for all bacterial samples. RNA samples were sent for dual RNA-sequencing (or bacterial RNA-sequencing for input controls), to the Microbial Genome Sequencing Center for Illumina Sequencing. Samples were rRNA depleted using RiboZero Plus (Illumina) and libraries for the 12 mouse samples and 3 input bacteria cDNA samples were sequenced using the Illumina Stranded RNA sequencing platform. Raw sequencing reads in fastq format were aligned and annotated to the mouse reference genome (mm10) or to the Clinical Group B Streptococcal Isolate CJB111 reference genome (CP063198_sRNA) using Qiagen CLC Genomics Workbench default settings (version 21.0.5): mismatch cost, 2; insertion and deletion cost, 3; length and similarity fraction, 0.8. Normalization and differential expression calculations of uniquely mapped mouse or bacterial transcripts were performed using R package DESeq2 (v1.34.0, RRID:SCR_015687) (78). Heatmaps and further comparison of gene expression changes for all mouse and bacterial samples were generated with Rstudio (v1.3.1073, RRID:SCR_000432). Gene set enrichment analysis (GSEA) was performed using the fGSEA R package (v1.20.0, RRID:SCR_020938) with 10,000 permutations and the Hallmarks and GO Biological Processes gene set collections from the Molecular Signatures Database (79).

### ELISAs on wound homogenates

Proteins in homogenized tissues was quantified using R&D systems ELISA kits (catalog # DY453, DY667, DY8596-05, DY4517-05). Protein detected was normalized to tissue weight and reported as protein (pg) per mg of tissue.

### Adherence of GBS to host cells

Adherence assays were performed as previously described (29, 80). Briefly, cell lines were seeded into 24 well plates and grown to a complete monolayer (approximately 1 × 10^6^ cells/well). GBS was grown to mid-log phase and normalized to 1 × 10^8^ CFU/mL in PBS. 1 × 10^6^ bacteria were added to one well of host cells to achieve a MOI of 1. To assess adherence of GBS to host cells, bacteria were incubated with host cells for 30 min then the cells were washed five times with PBS. Host cells were detached with 0.25% trypsin (Thermo Fisher Scientific) and permeabilized with 0.025% Triton X-100 (Sigma) in PBS, serially diluted, and plated to quantify all cell-associated bacteria.

### Statistical analyses

Statistical analysis was performed using Prism version 9.0.0 (121) for Windows (GraphPad Software, La Jolla, CA, United States) as described in figure legends.

## Supporting information

Supplemental Data Merged

## Acknowledgments

We would like to thank Elizabeth Grice and Sue Gardner for providing clinical GBS isolates, Laura Cook for providing the Δ*pbsP* mutant, Lakshmi Rajagopal for the Δ*covR* mutant, and Amber Nguyen for assistance with sequencing. This work was supported by NIH grants R01NS116716, R01AI153332 to K.S.D., NIHT32DK120521 to R.A.K. and R01AI153185 to A.R.H., and Department of Veteran Affairs Award BX002711 to A.R.H. The authors declare no other competing interests.

## Data and Materials Availability

All data needed to evaluate the conclusions in the paper are present in the paper and/or the Supplementary Materials. Mouse and bacterial RNA-sequencing datasets generated in this study including all raw data files were uploaded to NCBI GEO Depository (GSE201342).

## Supplemental figures and tables, and legends

**Supplemental Figure 1.**
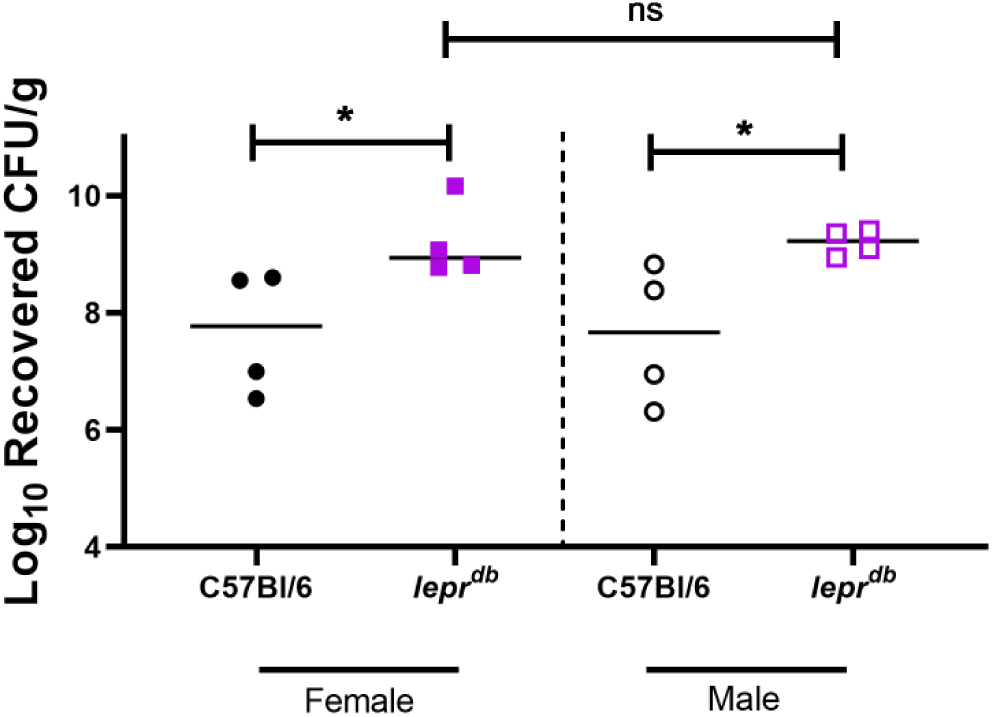
Murine Model of GBS Diabetic Wound Infection in Female and Male Mice. (A) CFU recovered from wounds of non-diabetic and diabetic mice after GBS infection. All animal infections proceeded for four days with three days under adhesive and sacrifice 24 h after adhesive removal. Significance determined by Mann–Whitney U test; *p<.05.

**Supplemental Table 1.**
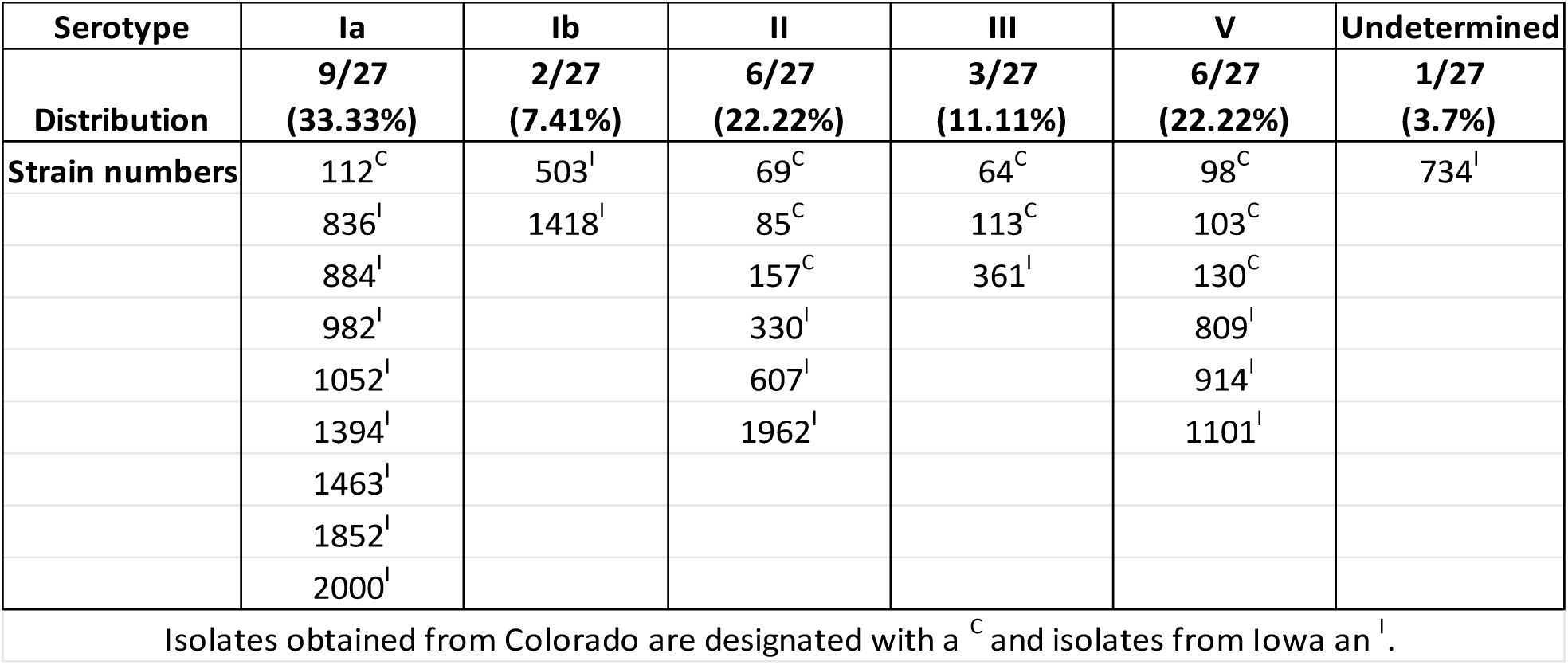
Clinical isolate serotypes.

**Supplemental Figure 2.**
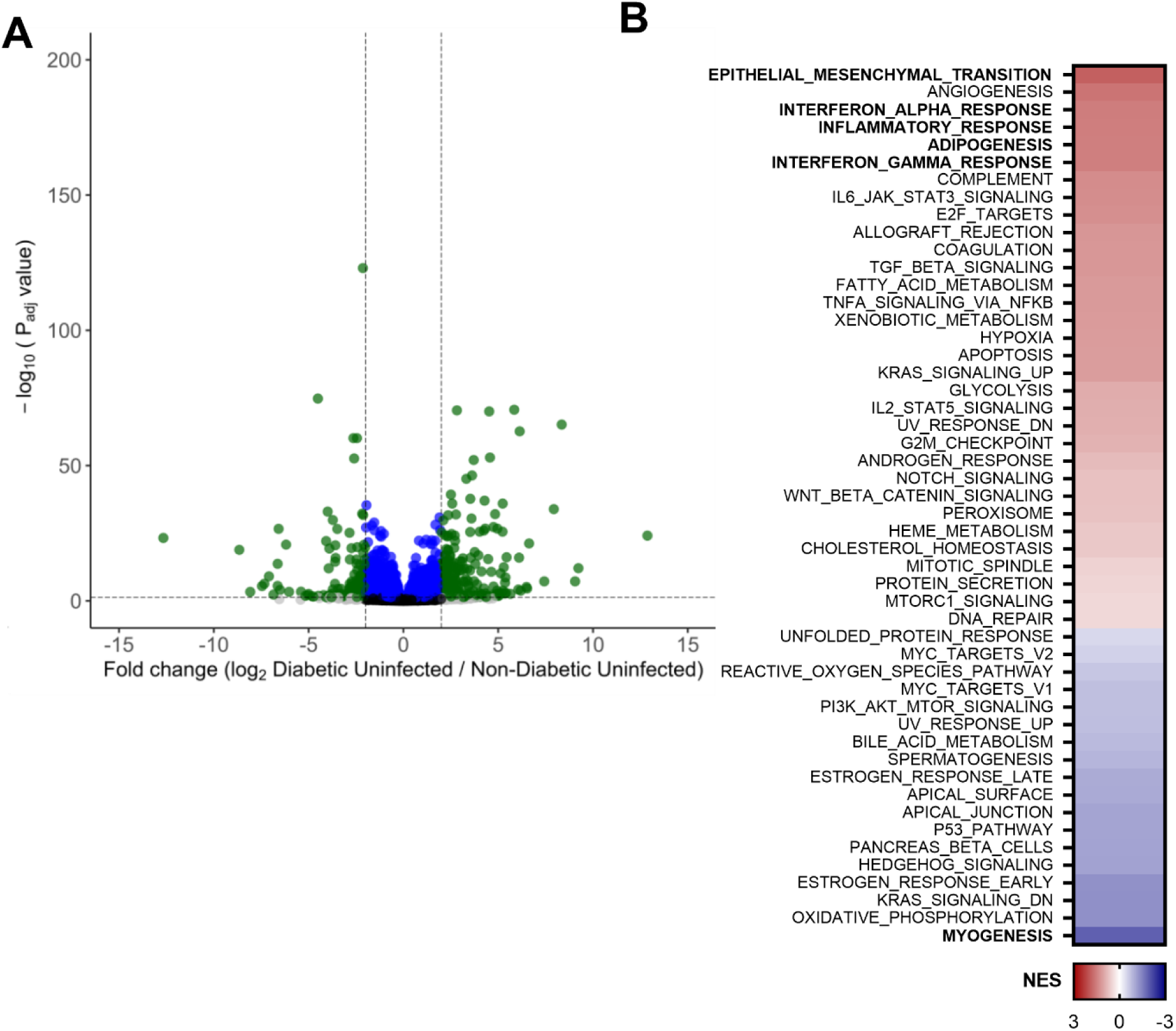
Murine transcriptome in non-diabetic uninfected vs. diabetic uninfected comparison. (A) Volcano plot of differentially expressed genes. (B) GSEA of pathways enriched in diabetic wounds. Significant pathways are in bold. NES presented as a heat map with highly enriched pathways in red.

**Supplemental Table 2.**
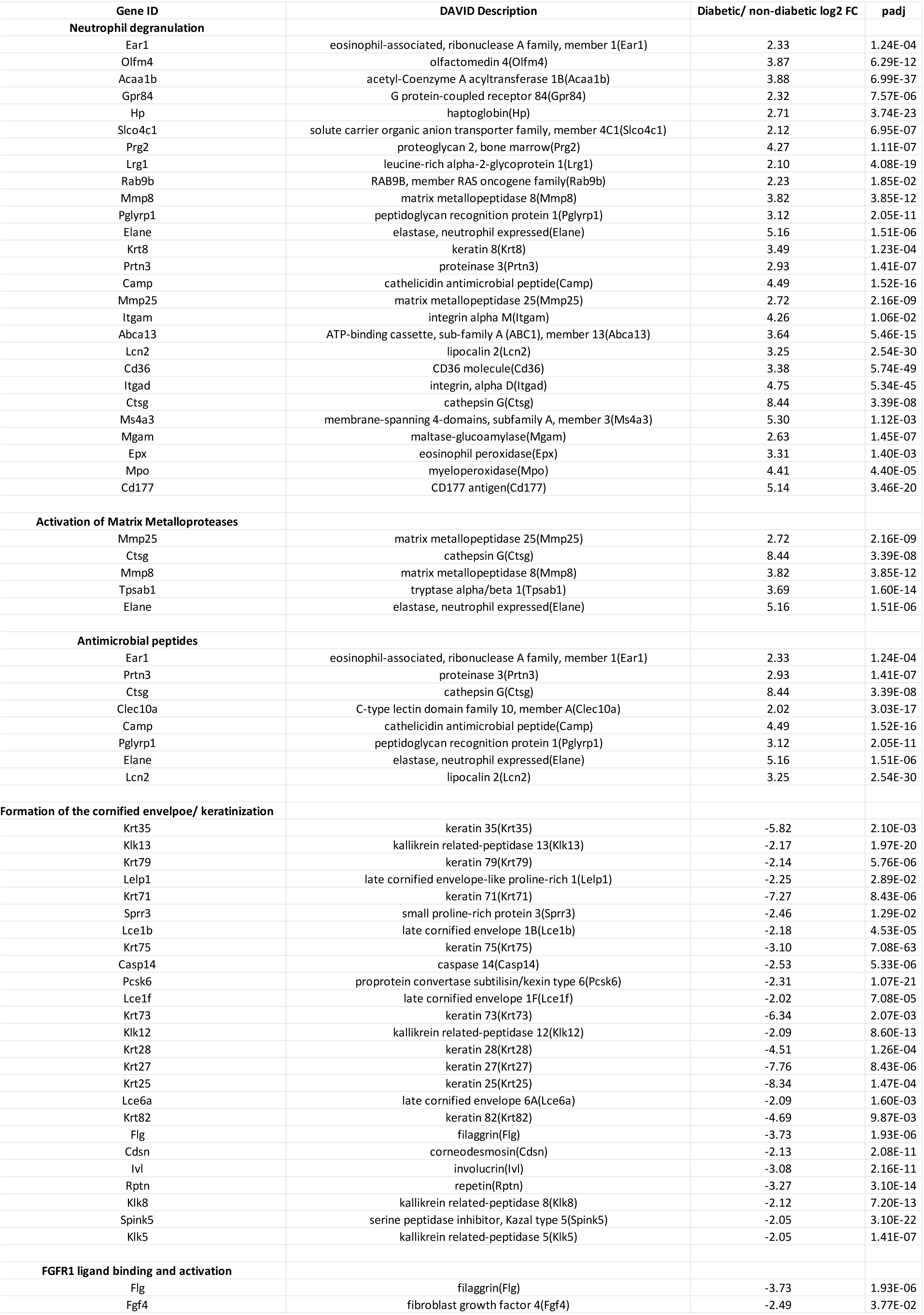
Murine transcriptome in diabetic infected vs. non-diabetic infected comparison. Select genes and pathways.

**Supplemental Table 3.**
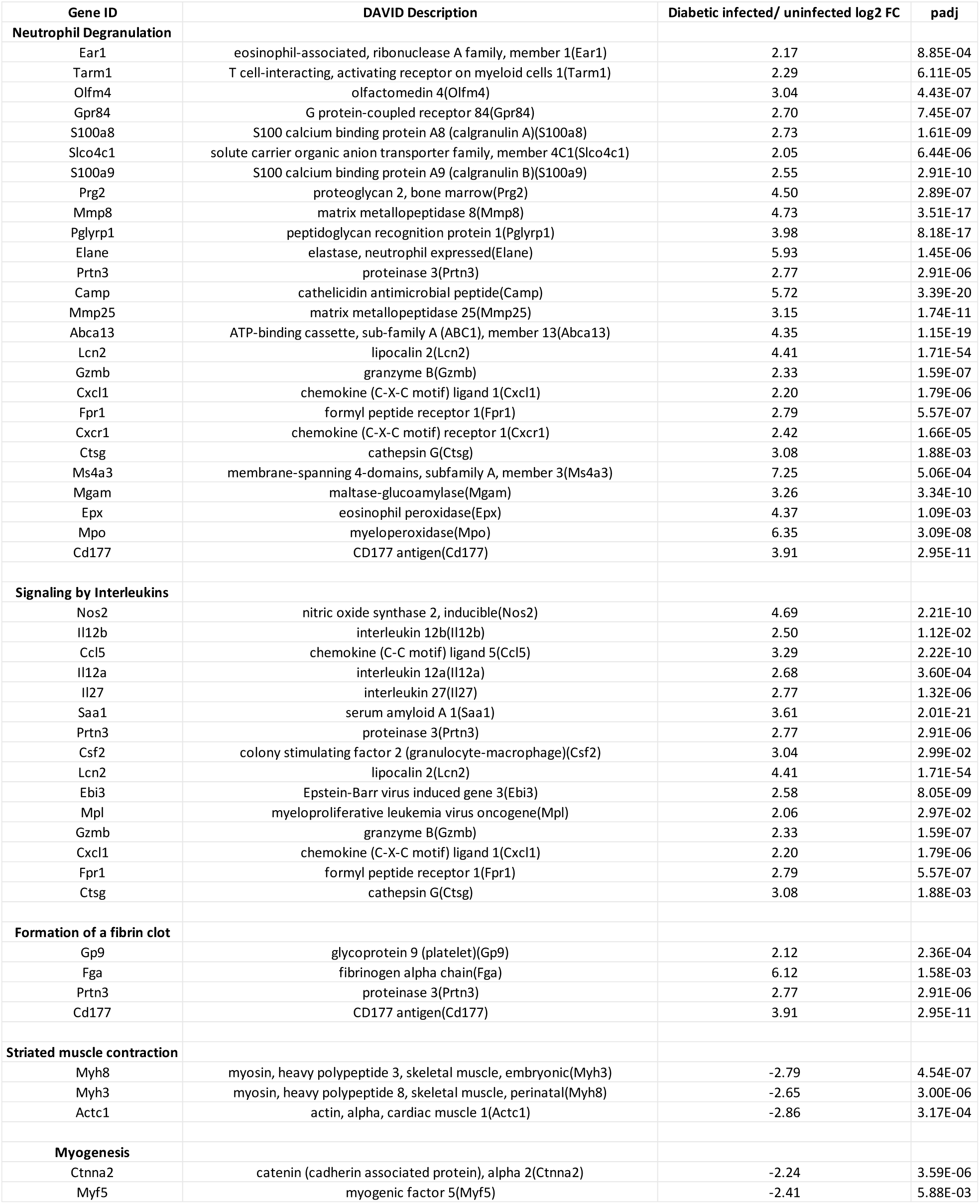
Murine transcriptome in diabetic infected vs. diabetic uninfected comparison. Select genes and pathways.

**Supplemental Table 4.**
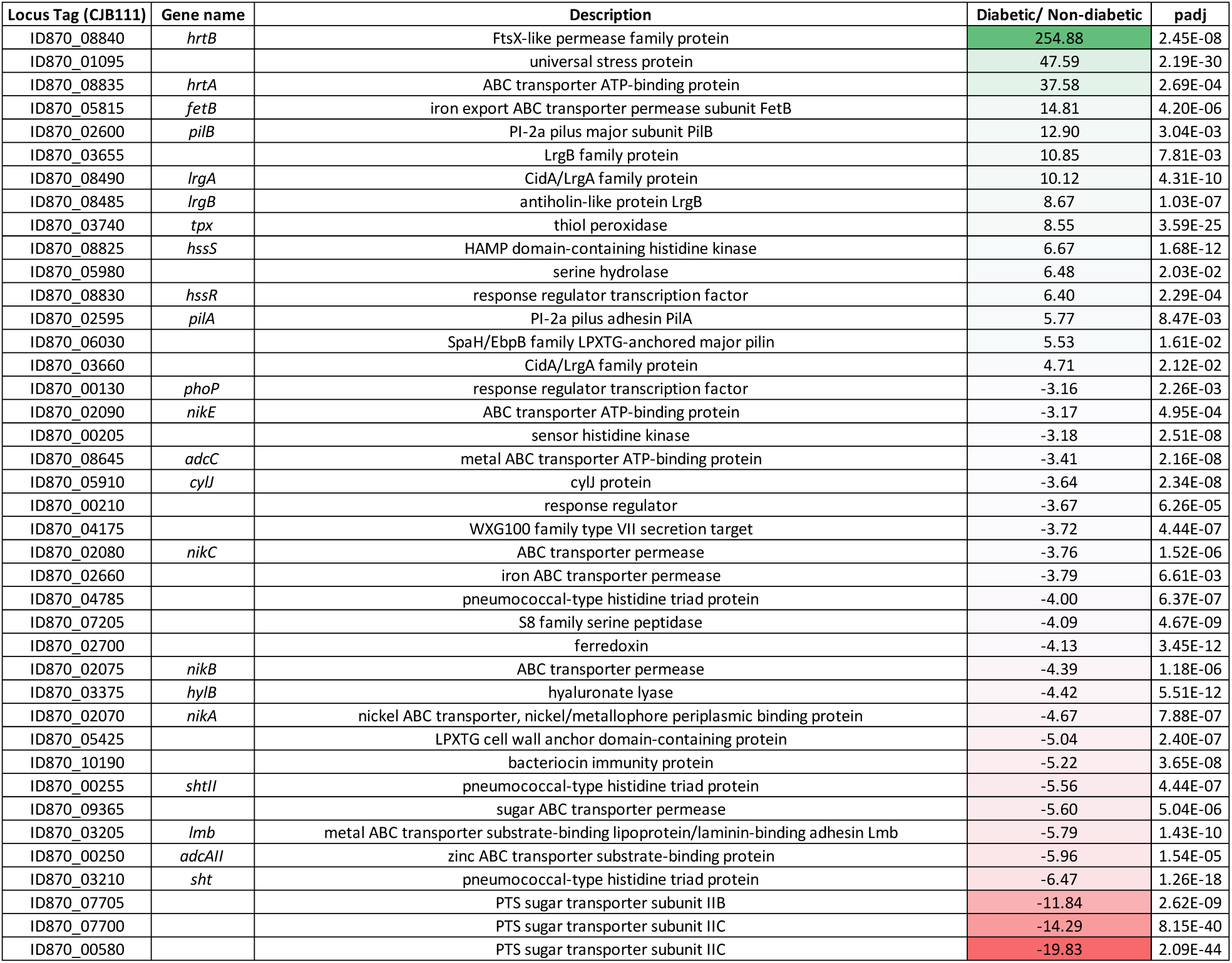
Select GBS transcripts with altered regulation in diabetic wounds vs. non-diabetic.

**Supplemental Table 5.**
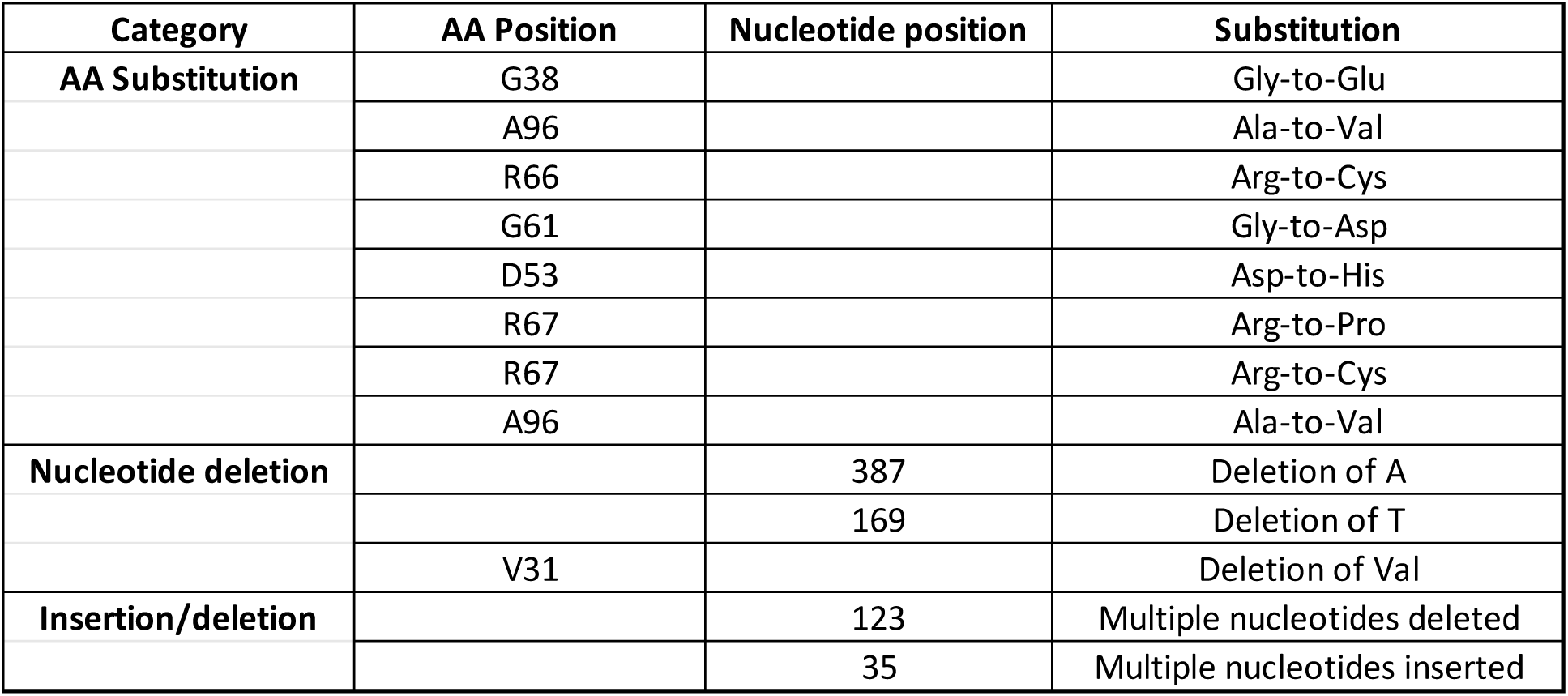
Mutations in *covR* from hyperpigmented colonies recovered from the diabetic wound.

**Supplemental Figure 3.**
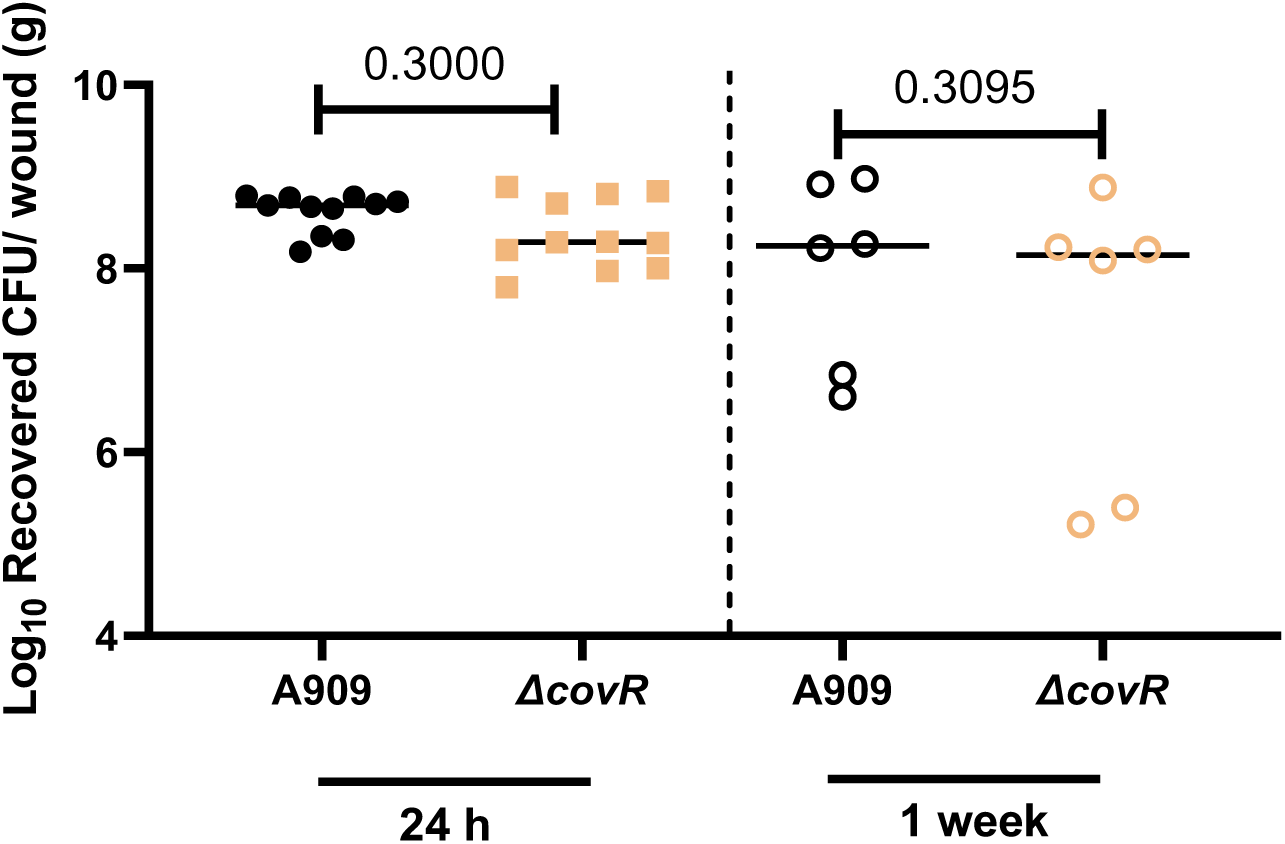
Analysis of a Δ*covR* mutant in diabetic wound infection. CFU recovered from wounds of diabetic mice after GBS infection. Animal infections proceeded for four or 11 days with three days under adhesive and sacrifice 24 h or 1 week after adhesive removal. Significance determined by Mann–Whitney U test.

**Supplemental Figure 4.**
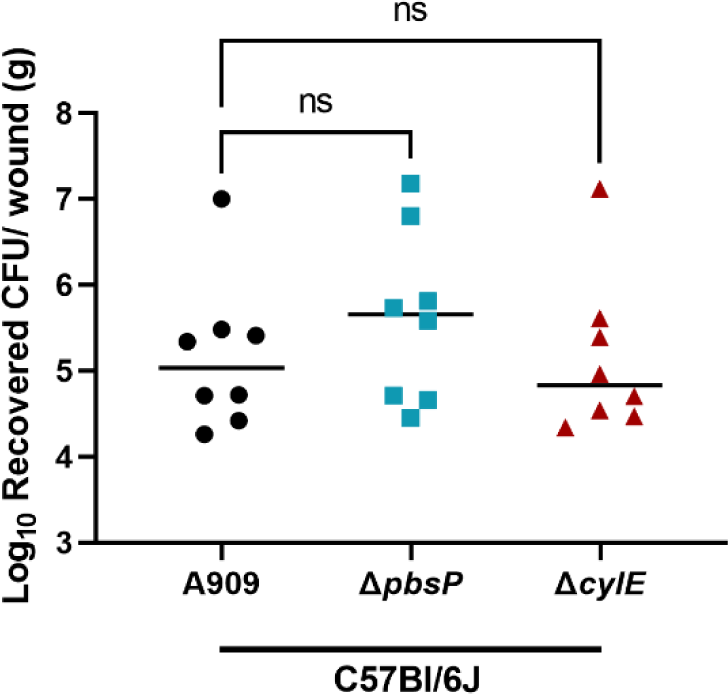
Analysis of a Δ*pbsP and* Δ*cylE* mutant in non-diabetic wound infection. CFU recovered from wounds of diabetic mice after GBS infection. Animal infections proceeded for four days with three days under adhesive and sacrifice 24 h after adhesive removal. Significance determined by One-way ANOVA with comparison to A909.

**Supplemental Table 6.**
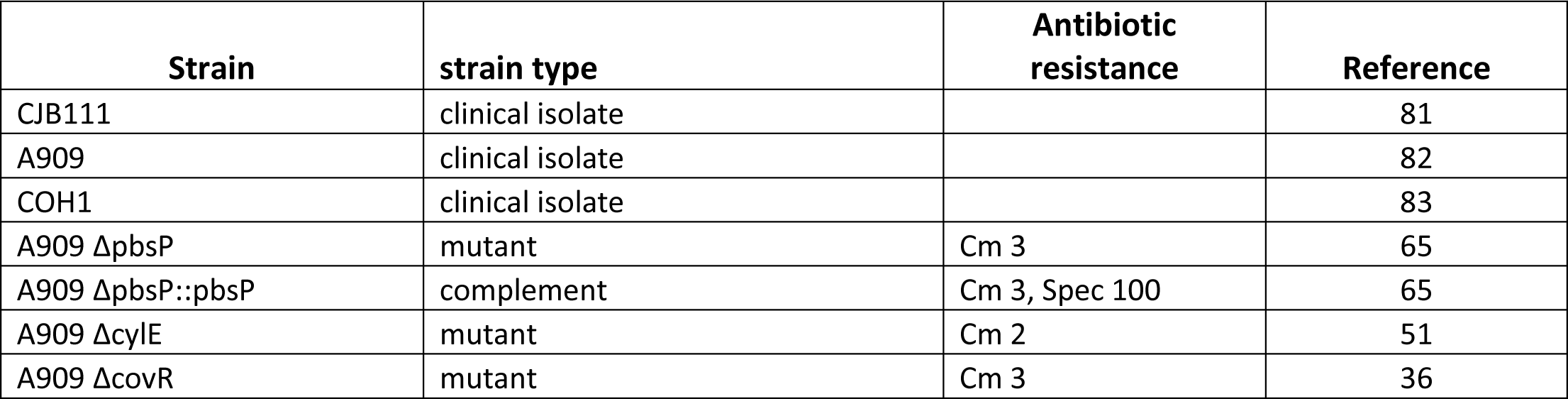
Strains used in this study

| Strain | strain type | Antibiotic resistance | Reference |
| --- | --- | --- | --- |
| CJB111 | clinical isolate |  | 81 |
| A909 | clinical isolate |  | 82 |
| COH1 | clinical isolate |  | 83 |
| A909 $\Delta pbsP$ | mutant | Cm 3 | 65 |
| A909 $\Delta pbsP::pbsP$ | complement | Cm 3, Spec 100 | 65 |
| A909 $\Delta cylE$ | mutant | Cm 2 | 51 |
| A909 $\Delta covR$ | mutant | Cm 3 | 36 |

**Supplemental Table 7.**
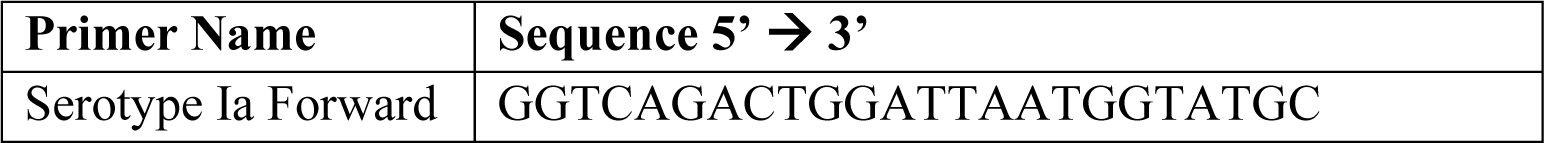
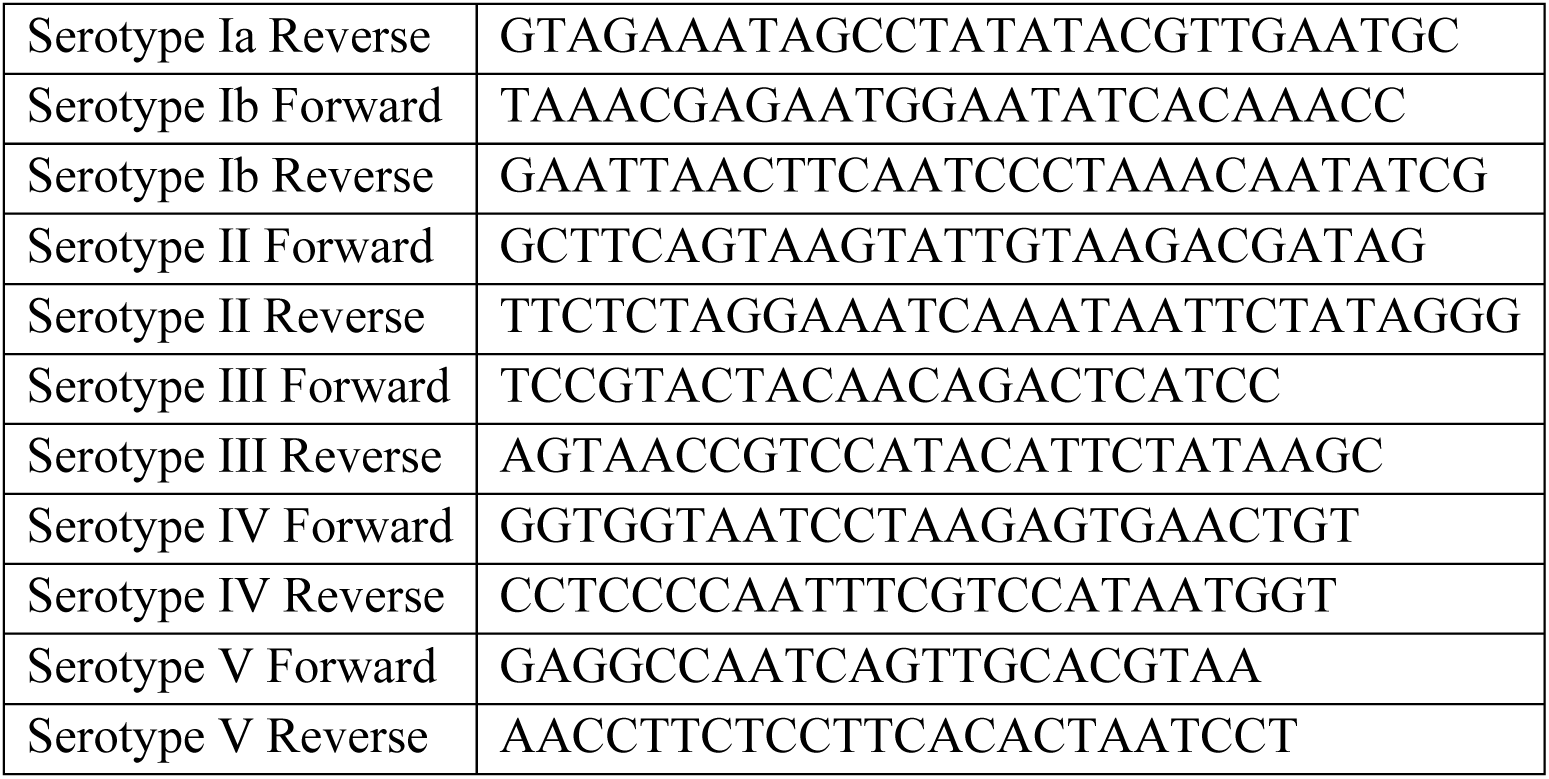
Primers used in this study

