## Supplemental Data Merged for "Group B Streptococcus Adaptation Promotes Survival in a Hyperinflammatory Diabetic Wound Environment"

Science Advances
AAAS

### Supplementary Materials for

#### Group B Streptococcus Adaptation Promotes Survival in a Hyperinflammatory Diabetic Wound Environment

Rebecca A. Keogh, Amanda L. Haeberle, Christophe J. Langouët-Astrié, Jeffrey S. Kavanaugh, Eric P. Schmidt, Garrett D. Moore, Alexander R. Horswill and Kelly S. Doran

**This PDF file includes:**

Figs. S1 to S4
Tables S1 to S7

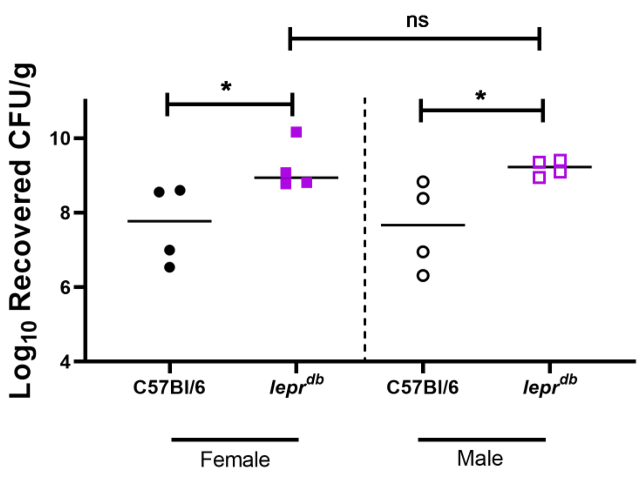

**Fig. S1. Murine Model of GBS Diabetic Wound Infection in Female and Male Mice.** (A) CFU recovered from wounds of non-diabetic and diabetic mice after GBS infection. All animal infections proceeded for four days with three days under adhesive and sacrifice 24 h after adhesive removal. Significance determined by Mann–Whitney U test; *p<.05.

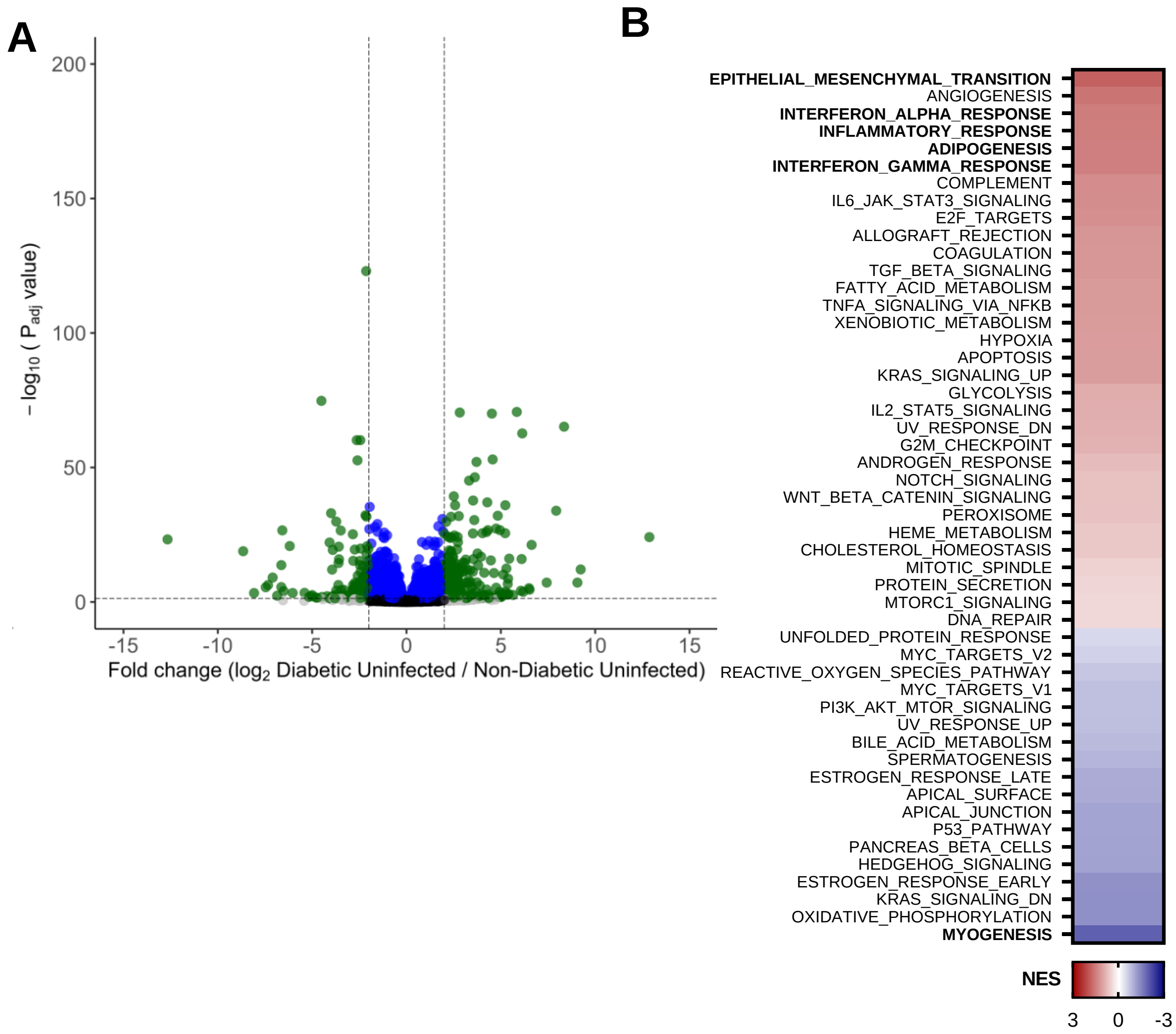

**Fig. S2.** Murine transcriptome in non-diabetic uninfected vs. diabetic uninfected comparison. (A) Volcano plot of differentially expressed genes. (B) GSEA of pathways enriched in diabetic wounds. Significant pathways are in bold. NES presented as a heat map with highly enriched pathways in red.

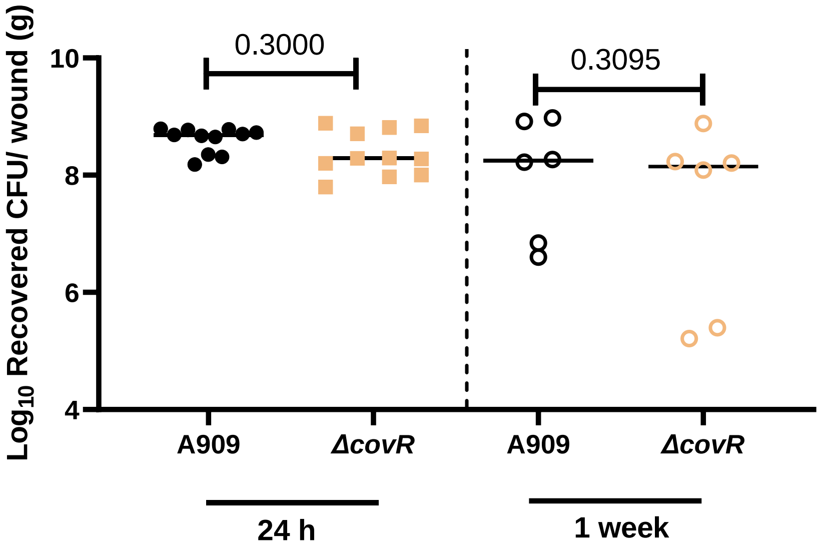

**Fig. S3.** Analysis of a Δ*covR* mutant in diabetic wound infection. CFU recovered from wounds of diabetic mice after GBS infection. Animal infections proceeded for four or 11 days with three days under adhesive and sacrifice 24 h or 1 week after adhesive removal. Significance determined by Mann–Whitney U test.

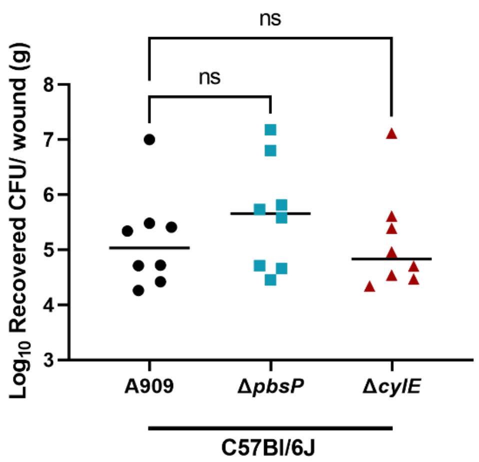

**Fig. S4.** Analysis of a Δ*pbsP and* Δ*cylE* mutant in non-diabetic wound infection. CFU recovered from wounds of diabetic mice after GBS infection. Animal infections proceeded for four days with three days under adhesive and sacrifice 24 h after adhesive removal. Significance determined by One-way ANOVA with comparison to A909.

| Serotype | Ia | Ib | II | III | V | Undetermined |
|---|---|---|---|---|---|---|
| Distribution | 9/27 (33.33%) | 2/27 (7.41%) | 6/27 (22.22%) | 3/27 (11.11%) | 6/27 (22.22%) | 1/27 (3.7%) |
| Strain numbers | 112[C] | 503[I] | 69[C] | 64[C] | 98[C] | 734[I] |
| | 836[I] | 1418[I] | 85[C] | 113[C] | 103[C] | |
| | 884[I] | | 157[C] | 361[I] | 130[C] | |
| | 982[I] | | 330[I] | | 809[I] | |
| | 1052[I] | | 607[I] | | 914[I] | |
| | 1394[I] | | 1962[I] | | 1101[I] | |
| | 1463[I] | | | | | |
| | 1852[I] | | | | | |
| | 2000[I] | | | | | |

Isolates obtained from Colorado are designated with a [C] and isolates from Iowa an [I].

**Table S1.** Clinical isolate serotypes.

| Gene ID | DAVID Description | Diabetic/ non-diabetic log2 FC | padj |
|---|---|---|---|
| **Neutrophil degranulation** | | | |
| Ear1 | eosinophil-associated, ribonuclease A family, member 1(Ear1) | 2.33 | 1.24E-04 |
| Olfm4 | olfactomedin 4(Olfm4) | 3.87 | 6.29E-12 |
| Acaa1b | acetyl-Coenzyme A acyltransferase 1B(Acaa1b) | 3.88 | 6.99E-37 |
| Gpr84 | G protein-coupled receptor 84(Gpr84) | 2.32 | 7.57E-06 |
| Hp | haptoglobin(Hp) | 2.71 | 3.74E-23 |
| Slco4c1 | solute carrier organic anion transporter family, member 4C1(Slco4c1) | 2.12 | 6.95E-07 |
| Prg2 | proteoglycan 2, bone marrow(Prg2) | 4.27 | 1.11E-07 |
| Lrg1 | leucine-rich alpha-2-glycoprotein 1(Lrg1) | 2.10 | 4.08E-19 |
| Rab9b | RAB9B, member RAS oncogene family(Rab9b) | 2.23 | 1.85E-02 |
| Mmp8 | matrix metallopeptidase 8(Mmp8) | 3.82 | 3.85E-12 |
| Pglyrp1 | peptidoglycan recognition protein 1(Pglyrp1) | 3.12 | 2.05E-11 |
| Elane | elastase, neutrophil expressed(Elane) | 5.16 | 1.51E-06 |
| Krt8 | keratin 8(Krt8) | 3.49 | 1.23E-04 |
| Prtn3 | proteinase 3(Prtn3) | 2.93 | 1.41E-07 |
| Camp | cathelicidin antimicrobial peptide(Camp) | 4.49 | 1.52E-16 |
| Mmp25 | matrix metallopeptidase 25(Mmp25) | 2.72 | 2.16E-09 |
| Itgam | integrin alpha M(Itgam) | 4.26 | 1.06E-02 |
| Abca13 | ATP-binding cassette, sub-family A (ABC1), member 13(Abca13) | 3.64 | 5.46E-15 |
| Lcn2 | lipocalin 2(Lcn2) | 3.25 | 2.54E-30 |
| Cd36 | CD36 molecule(Cd36) | 3.38 | 5.74E-49 |
| Itgad | integrin, alpha D(Itgad) | 4.75 | 5.34E-45 |
| Ctsg | cathepsin G(Ctsg) | 8.44 | 3.39E-08 |
| Ms4a3 | membrane-spanning 4-domains, subfamily A, member 3(Ms4a3) | 5.30 | 1.12E-03 |
| Mgam | maltase-glucoamylase(Mgam) | 2.63 | 1.45E-07 |
| Epx | eosinophil peroxidase(Epx) | 3.31 | 1.40E-03 |
| Mpo | myeloperoxidase(Mpo) | 4.41 | 4.40E-05 |
| Cd177 | CD177 antigen(Cd177) | 5.14 | 3.46E-20 |
| **Activation of Matrix Metalloproteases** | | | |
| Mmp25 | matrix metallopeptidase 25(Mmp25) | 2.72 | 2.16E-09 |
| Ctsg | cathepsin G(Ctsg) | 8.44 | 3.39E-08 |
| Mmp8 | matrix metallopeptidase 8(Mmp8) | 3.82 | 3.85E-12 |
| Tpsab1 | tryptase alpha/beta 1(Tpsab1) | 3.69 | 1.60E-14 |
| Elane | elastase, neutrophil expressed(Elane) | 5.16 | 1.51E-06 |
| **Antimicrobial peptides** | | | |
| Ear1 | eosinophil-associated, ribonuclease A family, member 1(Ear1) | 2.33 | 1.24E-04 |
| Prtn3 | proteinase 3(Prtn3) | 2.93 | 1.41E-07 |
| Ctsg | cathepsin G(Ctsg) | 8.44 | 3.39E-08 |
| Clec10a | C-type lectin domain family 10, member A(Clec10a) | 2.02 | 3.03E-17 |
| Camp | cathelicidin antimicrobial peptide(Camp) | 4.49 | 1.52E-16 |
| Pglyrp1 | peptidoglycan recognition protein 1(Pglyrp1) | 3.12 | 2.05E-11 |
| Elane | elastase, neutrophil expressed(Elane) | 5.16 | 1.51E-06 |
| Lcn2 | lipocalin 2(Lcn2) | 3.25 | 2.54E-30 |
| **Formation of the cornified envelpoe/ keratinization** | | | |
| Krt35 | keratin 35(Krt35) | -5.82 | 2.10E-03 |
| Klk13 | kallikrein related-peptidase 13(Klk13) | -2.17 | 1.97E-20 |
| Krt79 | keratin 79(Krt79) | -2.14 | 5.76E-06 |
| Lelp1 | late cornified envelope-like proline-rich 1(Lelp1) | -2.25 | 2.89E-02 |
| Krt71 | keratin 71(Krt71) | -7.27 | 8.43E-06 |
| Sprr3 | small proline-rich protein 3(Sprr3) | -2.46 | 1.29E-02 |
| Lce1b | late cornified envelope 1B(Lce1b) | -2.18 | 4.53E-05 |
| Krt75 | keratin 75(Krt75) | -3.10 | 7.08E-63 |
| Casp14 | caspase 14(Casp14) | -2.53 | 5.33E-06 |
| Pcsk6 | proprotein convertase subtilisin/kexin type 6(Pcsk6) | -2.31 | 1.07E-21 |
| Lce1f | late cornified envelope 1F(Lce1f) | -2.02 | 7.08E-05 |
| Krt73 | keratin 73(Krt73) | -6.34 | 2.07E-03 |
| Klk12 | kallikrein related-peptidase 12(Klk12) | -2.09 | 8.60E-13 |
| Krt28 | keratin 28(Krt28) | -4.51 | 1.26E-04 |
| Krt27 | keratin 27(Krt27) | -7.76 | 8.43E-06 |
| Krt25 | keratin 25(Krt25) | -8.34 | 1.47E-04 |
| Lce6a | late cornified envelope 6A(Lce6a) | -2.09 | 1.60E-03 |
| Krt82 | keratin 82(Krt82) | -4.69 | 9.87E-03 |
| Flg | filaggrin(Flg) | -3.73 | 1.93E-06 |
| Cdsn | corneodesmosin(Cdsn) | -2.13 | 2.08E-11 |
| Ivl | involucrin(Ivl) | -3.08 | 2.16E-11 |
| Rptn | repetin(Rptn) | -3.27 | 3.10E-14 |
| Klk8 | kallikrein related-peptidase 8(Klk8) | -2.12 | 7.20E-13 |
| Spink5 | serine peptidase inhibitor, Kazal type 5(Spink5) | -2.05 | 3.10E-22 |
| Klk5 | kallikrein related-peptidase 5(Klk5) | -2.05 | 1.41E-07 |
| **FGFR1 ligand binding and activation** | | | |
| Flg | filaggrin(Flg) | -3.73 | 1.93E-06 |
| Fgf4 | fibroblast growth factor 4(Fgf4) | -2.49 | 3.77E-02 |

**Table S2.** Murine transcriptome in diabetic infected vs. non-diabetic infected comparison. Select genes and pathways.

| Gene ID | DAVID Description | Diabetic infected/ uninfected log2 FC | padj |
|---|---|---|---|
| **Neutrophil Degranulation** | | | |
| Ear1 | eosinophil-associated, ribonuclease A family, member 1(Ear1) | 2.17 | 8.85E-04 |
| Tarm1 | T cell-interacting, activating receptor on myeloid cells 1(Tarm1) | 2.29 | 6.11E-05 |
| Olfm4 | olfactomedin 4(Olfm4) | 3.04 | 4.43E-07 |
| Gpr84 | G protein-coupled receptor 84(Gpr84) | 2.70 | 7.45E-07 |
| S100a8 | S100 calcium binding protein A8 (calgranulin A)(S100a8) | 2.73 | 1.61E-09 |
| Slco4c1 | solute carrier organic anion transporter family, member 4C1(Slco4c1) | 2.05 | 6.44E-06 |
| S100a9 | S100 calcium binding protein A9 (calgranulin B)(S100a9) | 2.55 | 2.91E-10 |
| Prg2 | proteoglycan 2, bone marrow(Prg2) | 4.50 | 2.89E-07 |
| Mmp8 | matrix metallopeptidase 8(Mmp8) | 4.73 | 3.51E-17 |
| Pglyrp1 | peptidoglycan recognition protein 1(Pglyrp1) | 3.98 | 8.18E-17 |
| Elane | elastase, neutrophil expressed(Elane) | 5.93 | 1.45E-06 |
| Prtn3 | proteinase 3(Prtn3) | 2.77 | 2.91E-06 |
| Camp | cathelicidin antimicrobial peptide(Camp) | 5.72 | 3.39E-20 |
| Mmp25 | matrix metallopeptidase 25(Mmp25) | 3.15 | 1.74E-11 |
| Abca13 | ATP-binding cassette, sub-family A (ABC1), member 13(Abca13) | 4.35 | 1.15E-19 |
| Lcn2 | lipocalin 2(Lcn2) | 4.41 | 1.71E-54 |
| Gzmb | granzyme B(Gzmb) | 2.33 | 1.59E-07 |
| Cxcl1 | chemokine (C-X-C motif) ligand 1(Cxcl1) | 2.20 | 1.79E-06 |
| Fpr1 | formyl peptide receptor 1(Fpr1) | 2.79 | 5.57E-07 |
| Cxcr1 | chemokine (C-X-C motif) receptor 1(Cxcr1) | 2.42 | 1.66E-05 |
| Ctsg | cathepsin G(Ctsg) | 3.08 | 1.88E-03 |
| Ms4a3 | membrane-spanning 4-domains, subfamily A, member 3(Ms4a3) | 7.25 | 5.06E-04 |
| Mgam | maltase-glucoamylase(Mgam) | 3.26 | 3.34E-10 |
| Epx | eosinophil peroxidase(Epx) | 4.37 | 1.09E-03 |
| Mpo | myeloperoxidase(Mpo) | 6.35 | 3.09E-08 |
| Cd177 | CD177 antigen(Cd177) | 3.91 | 2.95E-11 |
| **Signaling by Interleukins** | | | |
| Nos2 | nitric oxide synthase 2, inducible(Nos2) | 4.69 | 2.21E-10 |
| Il12b | interleukin 12b(Il12b) | 2.50 | 1.12E-02 |
| Ccl5 | chemokine (C-C motif) ligand 5(Ccl5) | 3.29 | 2.22E-10 |
| Il12a | interleukin 12a(Il12a) | 2.68 | 3.60E-04 |
| Il27 | interleukin 27(Il27) | 2.77 | 1.32E-06 |
| Saa1 | serum amyloid A 1(Saa1) | 3.61 | 2.01E-21 |
| Prtn3 | proteinase 3(Prtn3) | 2.77 | 2.91E-06 |
| Csf2 | colony stimulating factor 2 (granulocyte-macrophage)(Csf2) | 3.04 | 2.99E-02 |
| Lcn2 | lipocalin 2(Lcn2) | 4.41 | 1.71E-54 |
| Ebi3 | Epstein-Barr virus induced gene 3(Ebi3) | 2.58 | 8.05E-09 |
| Mpl | myeloproliferative leukemia virus oncogene(Mpl) | 2.06 | 2.97E-02 |
| Gzmb | granzyme B(Gzmb) | 2.33 | 1.59E-07 |
| Cxcl1 | chemokine (C-X-C motif) ligand 1(Cxcl1) | 2.20 | 1.79E-06 |
| Fpr1 | formyl peptide receptor 1(Fpr1) | 2.79 | 5.57E-07 |
| Ctsg | cathepsin G(Ctsg) | 3.08 | 1.88E-03 |
| **Formation of a fibrin clot** | | | |
| Gp9 | glycoprotein 9 (platelet)(Gp9) | 2.12 | 2.36E-04 |
| Fga | fibrinogen alpha chain(Fga) | 6.12 | 1.58E-03 |
| Prtn3 | proteinase 3(Prtn3) | 2.77 | 2.91E-06 |
| Cd177 | CD177 antigen(Cd177) | 3.91 | 2.95E-11 |
| **Striated muscle contraction** | | | |
| Myh8 | myosin, heavy polypeptide 3, skeletal muscle, embryonic(Myh3) | -2.79 | 4.54E-07 |
| Myh3 | myosin, heavy polypeptide 8, skeletal muscle, perinatal(Myh8) | -2.65 | 3.00E-06 |
| Actc1 | actin, alpha, cardiac muscle 1(Actc1) | -2.86 | 3.17E-04 |
| **Myogenesis** | | | |
| Ctnna2 | catenin (cadherin associated protein), alpha 2(Ctnna2) | -2.24 | 3.59E-06 |
| Myf5 | myogenic factor 5(Myf5) | -2.41 | 5.88E-03 |

**Table S3.** Murine transcriptome in diabetic infected vs. diabetic uninfected comparison. Select genes and pathways.

| Locus Tag (CJB111) | Gene name | Description | Diabetic/ Non-diabetic | padj |
|---|---|---|---|---|
| ID870_08840 | *hrtB* | FtsX-like permease family protein | 254.88 | 2.45E-08 |
| ID870_01095 | | universal stress protein | 47.59 | 2.19E-30 |
| ID870_08835 | *hrtA* | ABC transporter ATP-binding protein | 37.58 | 2.69E-04 |
| ID870_05815 | *fetB* | iron export ABC transporter permease subunit FetB | 14.81 | 4.20E-06 |
| ID870_02600 | *pilB* | PI-2a pilus major subunit PilB | 12.90 | 3.04E-03 |
| ID870_03655 | | LrgB family protein | 10.85 | 7.81E-03 |
| ID870_08490 | *lrgA* | CidA/LrgA family protein | 10.12 | 4.31E-10 |
| ID870_08485 | *lrgB* | antiholin-like protein LrgB | 8.67 | 1.03E-07 |
| ID870_03740 | *tpx* | thiol peroxidase | 8.55 | 3.59E-25 |
| ID870_08825 | *hssS* | HAMP domain-containing histidine kinase | 6.67 | 1.68E-12 |
| ID870_05980 | | serine hydrolase | 6.48 | 2.03E-02 |
| ID870_08830 | *hssR* | response regulator transcription factor | 6.40 | 2.29E-04 |
| ID870_02595 | *pilA* | PI-2a pilus adhesin PilA | 5.77 | 8.47E-03 |
| ID870_06030 | | SpaH/EbpB family LPXTG-anchored major pilin | 5.53 | 1.61E-02 |
| ID870_03660 | | CidA/LrgA family protein | 4.71 | 2.12E-02 |
| ID870_00130 | *phoP* | response regulator transcription factor | -3.16 | 2.26E-03 |
| ID870_02090 | *nikE* | ABC transporter ATP-binding protein | -3.17 | 4.95E-04 |
| ID870_00205 | | sensor histidine kinase | -3.18 | 2.51E-08 |
| ID870_08645 | *adcC* | metal ABC transporter ATP-binding protein | -3.41 | 2.16E-08 |
| ID870_05910 | *cylJ* | cylJ protein | -3.64 | 2.34E-08 |
| ID870_00210 | | response regulator | -3.67 | 6.26E-05 |
| ID870_04175 | | WXG100 family type VII secretion target | -3.72 | 4.44E-07 |
| ID870_02080 | *nikC* | ABC transporter permease | -3.76 | 1.52E-06 |
| ID870_02660 | | iron ABC transporter permease | -3.79 | 6.61E-03 |
| ID870_04785 | | pneumococcal-type histidine triad protein | -4.00 | 6.37E-07 |
| ID870_07205 | | S8 family serine peptidase | -4.09 | 4.67E-09 |
| ID870_02700 | | ferredoxin | -4.13 | 3.45E-12 |
| ID870_02075 | *nikB* | ABC transporter permease | -4.39 | 1.18E-06 |
| ID870_03375 | *hylB* | hyaluronate lyase | -4.42 | 5.51E-12 |
| ID870_02070 | *nikA* | nickel ABC transporter, nickel/metallophore periplasmic binding protein | -4.67 | 7.88E-07 |
| ID870_05425 | | LPXTG cell wall anchor domain-containing protein | -5.04 | 2.40E-07 |
| ID870_10190 | | bacteriocin immunity protein | -5.22 | 3.65E-08 |
| ID870_00255 | *shtII* | pneumococcal-type histidine triad protein | -5.56 | 4.44E-07 |
| ID870_09365 | | sugar ABC transporter permease | -5.60 | 5.04E-06 |
| ID870_03205 | *lmb* | metal ABC transporter substrate-binding lipoprotein/laminin-binding adhesin Lmb | -5.79 | 1.43E-10 |
| ID870_00250 | *adcAII* | zinc ABC transporter substrate-binding protein | -5.96 | 1.54E-05 |
| ID870_03210 | *sht* | pneumococcal-type histidine triad protein | -6.47 | 1.26E-18 |
| ID870_07705 | | PTS sugar transporter subunit IIB | -11.84 | 2.62E-09 |
| ID870_07700 | | PTS sugar transporter subunit IIC | -14.29 | 8.15E-40 |
| ID870_00580 | | PTS sugar transporter subunit IIC | -19.83 | 2.09E-44 |

**Table S4**. Select GBS transcripts with altered regulation in diabetic wounds vs. non-diabetic.

| Category | AA Position | Nucleotide position | Substitution |
|---|---|---|---|
| **AA Substitution** | G38 | | Gly-to-Glu |
| | A96 | | Ala-to-Val |
| | R66 | | Arg-to-Cys |
| | G61 | | Gly-to-Asp |
| | D53 | | Asp-to-His |
| | R67 | | Arg-to-Pro |
| | R67 | | Arg-to-Cys |
| | A96 | | Ala-to-Val |
| **Nucleotide deletion** | | 387 | Deletion of A |
| | | 169 | Deletion of T |
| | V31 | | Deletion of Val |
| **Insertion/deletion** | | 123 | Multiple nucleotides deleted |
| | | 35 | Multiple nucleotides inserted |

**Table S5**. Select GBS transcripts with altered regulation in diabetic wounds vs. non-diabetic.

| Strain | strain type | Antibiotic resistance | Reference |
|---|---|---|---|
| CJB111 | clinical isolate | | 81 |
| A909 | clinical isolate | | 82 |
| COH1 | clinical isolate | | 83 |
| A909 ΔpbsP | mutant | Cm 3 | 65 |
| A909 ΔpbsP::pbsP | complement | Cm 3, Spec 100 | 65 |
| A909 ΔcylE | mutant | Cm 2 | 51 |
| A909 ΔcovR | mutant | Cm 3 | 36 |

**Table S6**. Strains used in this study.

| Primer Name | Sequence 5’ → 3’ |
|---|---|
| Serotype Ia Forward | GGTCAGACTGGATTAATGGTATGC |
| Serotype Ia Reverse | GTAGAAATAGCCTATATACGTTGAATGC |
| Serotype Ib Forward | TAAACGAGAATGGAATATCACAAACC |
| Serotype Ib Reverse | GAATTAACTTCAATCCCTAAACAATATCG |
| Serotype II Forward | GCTTCAGTAAGTATTGTAAGACGATAG |
| Serotype II Reverse | TTCTCTAGGAAATCAAATAATTCTATAGGG |
| Serotype III Forward | TCCGTACTACAACAGACTCATCC |
| Serotype III Reverse | AGTAACCGTCCATACATTCTATAAGC |
| Serotype IV Forward | GGTGGTAATCCTAAGAGTGAACTGT |
| Serotype IV Reverse | CCTCCCCAATTTCGTCCATAATGGT |
| Serotype V Forward | GAGGCCAATCAGTTGCACGTAA |
| Serotype V Reverse | AACCTTCTCCTTCACACTAATCCT |

**Table S7**. Primers used in this study.
